# CyanoGate: A Golden Gate modular cloning suite for engineering cyanobacteria based on the plant MoClo syntax

**DOI:** 10.1101/426700

**Authors:** Ravendran Vasudevan, Grant A.R. Gale, Alejandra A. Schiavon, Anton Puzorjov, John Malm, Michael D. Gillespie, Konstantinos Vavitsas, Valentin Zulkower, Baojun Wang, Christopher J. Howe, David Lea-Smith, Alistair J. McCormick

## Abstract

Recent advances in synthetic biology research have been underpinned by an exponential increase in available genomic information and a proliferation of advanced DNA assembly tools. The adoption of plasmid vector assembly standards and parts libraries has greatly enhanced the reproducibility of research and exchange of parts between different labs and biological systems. However, a standardised Modular Cloning (MoClo) system is not yet available for cyanobacteria, which lag behind other prokaryotes in synthetic biology despite their huge potential in biotechnological applications. By building on the assembly library and syntax of the Plant Golden Gate MoClo kit, we have developed a versatile system called CyanoGate that unites cyanobacteria with plant and algal systems. We have generated a suite of parts and acceptor vectors for making i) marked/unmarked knock-outs or integrations using an integrative acceptor vector, and ii) transient multigene expression and repression systems using known and novel replicative vectors. We have tested and compared the CyanoGate system in the established model cyanobacterium *Synechocystis* sp. PCC 6803 and the more recently described fast-growing strain *Synechococcus elongatus* UTEX 2973. The system is publicly available and can be readily expanded to accommodate other standardised MoClo parts.

## INTRODUCTION

Much work is focused on expanding synthetic biology approaches to engineer photosynthetic organisms, including cyanobacteria. Cyanobacteria are an evolutionarily ancient and diverse phylum of photosynthetic prokaryotic organisms that are ecologically important, and are thought to contribute *ca*. 25% to oceanic net primary productivity (1, 2). The chloroplasts of all photosynthetic eukaryotes, including plants, resulted from the endosymbiotic uptake of a cyanobacterium by a eukaryotic ancestor (3). Therefore, cyanobacteria have proved useful as model organisms for the study of photosynthesis, electron transport and associated biochemical pathways, many of which are conserved in eukaryotic algae and higher plants. Several unique aspects of cyanobacterial photosynthesis, such as the biophysical carbon concentrating mechanism, also show promise as a means for enhancing productivity in crop plants (4). Furthermore, cyanobacteria are increasingly recognized as valuable platforms for industrial biotechnology to convert CO_2_ and H_2_O into valuable products using solar energy (5-7). They are metabolically diverse and encode many components (e.g. P450 cytochromes) necessary for generating high-value pharmaceutical products that can be challenging to produce in other systems (8-11). Furthermore, cyanobacteria show significant promise in biophotovoltaic devices for generating electrical energy (12, 13).

Based on morphological complexity, cyanobacteria are classified into five sub-sections (I-V) (1). Several members of the five sub-sections have been reportedly transformed (14,15), suggesting that many cyanobacterial species are amenable to genetic manipulation. Exogenous DNA can be integrated into or removed from the genome through homologous recombination-based approaches using natural transformation, conjugation (tri-parental mating), or electroporation (16). Exogenous DNA can also be propagated by replicative vectors, although the latter are currently restricted to the use of a single vector-type based on the broad-host range RSF1010 origin (17-19). Transformation tools have been developed for generating “unmarked” mutant strains (lacking an antibiotic resistance marker cassette) in several model species, such as *Synechocystis* sp. PCC 6803 (*Synechocystis* hereafter) (20). More recently, markerless genome editing using CRISPR-based approaches has been demonstrated to function in both unicellular and filamentous strains (21,22).

Although exciting progress is being made in developing effective transformation systems, cyanobacteria still lag behind in the field of synthetic biology compared to bacterial (heterotrophic), yeast and mammalian systems. Relatively few broad-host range genetic parts have been characterised, but many libraries of parts for constructing regulatory modules and circuits are starting to become available, albeit using different standards, which makes them difficult to combine (23-34). One key challenge is clear: parts that are widely used in *Escherichia coli* behave very differently in model cyanobacterial species, such as *Synechocystis* (16). Furthermore, different cyanobacterial strains generally show a wide variation for functionality and performance of different genetic parts (e.g. promoters, reporter genes and antibiotic resistance markers (19, 27, 28, 30). This suggests that parts need to be validated, calibrated, and perhaps modified, for individual strains, including model species and strains that may be more commercially relevant. Rapid cloning and assembly methods are essential for accelerating the ‘design, build, test and learn’ cycle, which is a central tenet of synthetic biology (35).

The adoption of vector assembly standards and part libraries has greatly enhanced the scalability of synthetic biology-based approaches in a range of biological systems (36). Recent advances in synthetic biology have led to the development of standards for Type IIS restriction endonuclease-mediated assembly (commonly known as Golden Gate cloning) for several model systems, including plants (37-39). Based on a common Golden Gate Modular Cloning (MoClo) syntax, large libraries are now available for fusion of different genetic parts to assemble complex vectors cheaply and easily without proprietary tools and reagents (40). High-throughput and automated assembly are projected to be widely available soon through DNA synthesis and construction facilities, such as the UK DNA Synthesis Foundries, where MoClo is seen as the most suitable assembly standard (41).

Here we have developed an easy-to-use system called CyanoGate that unites cyanobacteria with plant and algal systems, by building on the established Golden Gate MoClo syntax and assembly library for plants (38) that has been adopted by the OpenPlant consortium (www.openplant.org), iGEM competitions as “Phytobricks” and the MoClo kit for the microalga *Chlamydomonas reinhardtii* (42). Firstly, we have constructed and characterised a suite of known and novel genetic parts (level 0) for use in cyanobacterial research, including promoters, terminators, antibiotic resistant markers, neutral sites and gene repression systems (43-45). Secondly, we have designed an additional level of acceptor vectors (level T) to facilitate integrative or replicative transformation. We have characterised assembled level T vectors in *Synechocystis* and in *Synechococcus elongatus* UTEX 2973 (UTEX 2973 hereafter), which has a reported doubling time similar to that of *Saccharomyces cerevisiae* under specific growth conditions (46, 47). Lastly, we have developed an online tool for assembly of CyanoGate and Plant MoClo vectors to assist with the adoption of the CyanoGate system for developing synthetic biology approaches in other cyanobacterial strains.

## MATERIALS AND METHODS

### Cyanobacterial culture conditions

Cyanobacterial strains of *Synechocystis*, UTEX 2973 and *Synechococcus elongatus* PCC 7942 (PCC 7942 hereafter) were maintained on 1.5% (w/v) agar plates containing BG11 medium. Liquid cultures were grown in Erlenmeyer flasks (100 ml) containing BG11 medium (48) supplemented with 10 mM NaHCO_3_, shaken at 100 rpm and aerated with filter-sterilised water-saturated atmospheric air. *Synechocystis* and PCC 7942 strains were grown at 30°C with continuous light (100 μmol photons m^−2^ s^−1^) and UTEX 2793 strains were grown at 40°C with 300 μmol photons m^−2^ s^−1^ in an Infors Multitron-Pro supplied with warm white LED lighting (Infors HT).

### Growth analysis

Growth of *Synechocystis*, UTEX 2973 and PCC 7942 was measured in a Photon Systems Instrument Multicultivator MC 1000-0D (MC). Starter cultures were grown in a Photon Systems Instrument AlgaeTron AG 230 at 30°C under continuous warm-white light (100 μmol photons m^−2^ s^−1^) with air bubbling and shaken at 160 rpm unless otherwise indicated. These were grown to an optical density at 750 nm (OD_750_) of approximately 1.0, and used to seed 80 ml cultures for growth in the MC at a starting OD_720_ of ~0.2. Cultures were then grown under continuous warm-white light at 30°C (300 μmol photons m^−2^ s^−1^) with air bubbling or 30°C (under 300 or 500 μmol photons m^−2^ s^−1^) or 41, 45 and 50°C (500 μmol photons m^−2^ s^−1^) with 5% CO_2_ bubbling until the fastest grown strain was at OD_720_ = ~0.9, the maximum accurate OD that can be measured with this device. A total of 5-6 replicate experiments were performed over two separate runs (16 in total), each inoculated from a different starter culture.

### Fluorescence assays

Transgenic strains maintained on agar plates containing appropriate antibiotics were used to inoculate 10 ml seed cultures that were grown to an optical density at 750 nm (OD_750_) of approximately 1.0, as measured with a WPA Biowave II spectrometer (Biochrom). Seed cultures were diluted to an OD750 of 0.2, and 2 ml starting cultures were transferred to 24-well plates (Costar^^®^^ Corning Incorporated) for experiments. *Synechocystis* and UTEX 2973 strains were grown in the same culturing conditions described above. OD_750_ was measured using a FLUOstar OMEGA microplate reader (BMG Labtech). Fluorescence of eYFP for individual cells (10,000 cells per culture) was measured by flow cytometry using an Attune NxT Flow Cytometer (Thermofisher). Cells were gated using forward and side scatter, and median eYFP fluorescence was calculated from excitation/emission wavelengths 488 nm/515-545 nm (49) and reported at 48 hr unless otherwise stated.

### Vector construction

#### Level 0 vectors

Native cyanobacterial genetic parts were amplified from genomic DNA using NEB Q5 High-Fidelity DNA Polymerase (New England Biolabs) (Fig. 1; Supp. Info. 1). Where necessary, native genetic parts were domesticated (i.e. *Bsa*I and *Bpi*I sites were removed) using specific primers. Alternatively, parts were synthesised as Gblocks^^®^^ DNA fragments (Integrated DNA Technology) and cloned directly into an appropriate level 0 acceptor (see Supp. Info. 2 for vector maps) (38).

**Figure 1.**
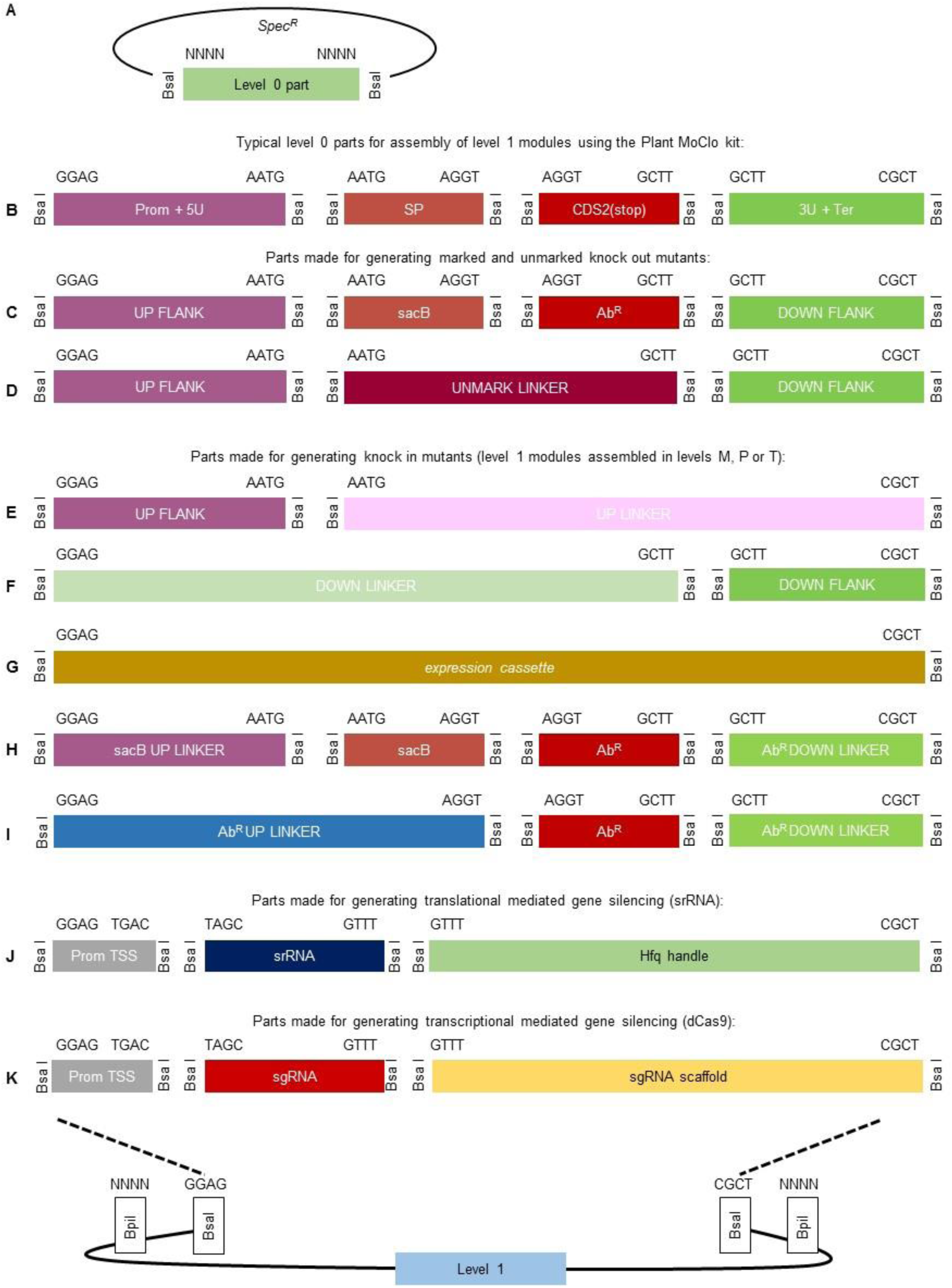
Adaptation of the Plant Golden Gate MoClo level 0 syntax for generating level 1 assemblies for transfer to Level T. (A) The format for a level 0 MoClo acceptor vector with the part bordered by two Bsal sites. (B) Typical level 0 parts from the Plant MoClo kit (38), where parts of the same type are bordered by the same pair of fusion sites (for each fusion site, only the sequence of the top strand is shown). Note that the parts are not drawn to scale. (C, D) The syntax of the Plant MoClo kit was adapted to generate level 0 parts for engineering marked and unmarked cyanobacterial mutant strains (20). (E-I) To generate knock in mutants, short linker parts (30 bp) were constructed to allow assembly of individual flanking sequences, or marker cassettes (Ab*^R^* or *sac*B) in level 1 vectors for subsequent assembly in level T. (J, K) Parts required for generating synthetic srRNA or CRISPRi level 1 constructs. See **Supp. Info 4 and 5** for workflows. Abbreviations: 3U+Ter, 3’UTR and terminator; Ab*^R^*, antibiotic resistance cassette; Ab*^R^* DOWN LINKER, short sequence (~30 bp) to provide CGCT overhang; Ab*^R^* UP LINKER, short sequence (~30 bp) to provide GAGG overhang; CDS2(stop), coding sequence with a stop codon; DOWN FLANK, flanking sequence downstream of target site; DOWN FLANK LINKER, short sequence (~30 bp) to provide GGAG overhang; Prom+5U, promoter and 5’ UTR; Prom TSS, promoter transcription start site; *sac*B, levansucrase expression cassette; *sac*B UP LINKER, short sequence (~30 bp) to provide GAGG overhang; sgRNA, single guide RNA; SP, signal peptide; srRNA, small regulatory RNA; UP FLANK, flanking sequence upstream of target site; UP FLANK LINKER, short sequence (~30 bp) to provide CGCT overhang; UNMARK LINKER, short sequence to bridge UP FLANK and DOWN FLANK.

Golden Gate assembly reactions were performed with restriction enzymes *Bsa*I (New England Biolabs) or *Bpi*I (Thermofisher), and T4 DNA ligase (Thermofisher) (see Supp. Info. 3-5 for detailed protocols). Vectors were transformed into One Shot TOP10 chemically competent *Escherichia coli* (Thermofisher) as per the manufacturer’s instructions. Transformed cultures were grown at 37°C on [1.5% (w/v)] LB agar plates or in liquid LB medium shaking at 260 rpm, with appropriate antibiotic selection for level 0, 1, M and P vectors as outlined in Engler et al. (38).

#### Level T acceptor vectors and new level 0 acceptors

A new level T vector system was designed that provides MoClo-compatible replicative vectors or integrative vectors for genomic modifications in cyanobacteria (16) (Fig. 2; Supp. Info. 1; 2). For replicative vectors, we modified the pPMQAK1 carrying an RSF1010 replicative origin (18) to make pPMQAK1-T, and vector pSEVA421 from the Standard European Vector Architecture (SEVA) 2.0 database (seva.cnb.csic.es) carrying the RK2 replicative origin to make pSEVA421-T (50). Replicative vector backbones were domesticated to remove native *Bsa*I and *Bpi*I sites where appropriate. The region between the BioBrick’s prefix and suffix was then replaced by a *lacZ* expression cassette flanked by two *Bpi*I sites that produce overhangs TGCC and GGGA, which are compatible with the plant Golden Gate MoClo assembly syntax for level 2 acceptors (e.g. pAGM4673) (38). For integrative vectors, we domesticated a pUC19 vector backbone and introduced two *Bpi*I sites compatible with a level 2 acceptor (as above) to make pUC19A-T and pUC19S-T. In addition, we made a new low copy level 0 acceptor (pSC101 origin of replication) for promoter parts based on the BioBrick standard vector pSB4K5 (51). DNA was amplified using NEB Q5 High-Fidelity DNA Polymerase (New England Biolabs). All vectors were sequenced following assembly to confirm domestication and the integrity of the MoClo cloning site.

**Figure 2.**
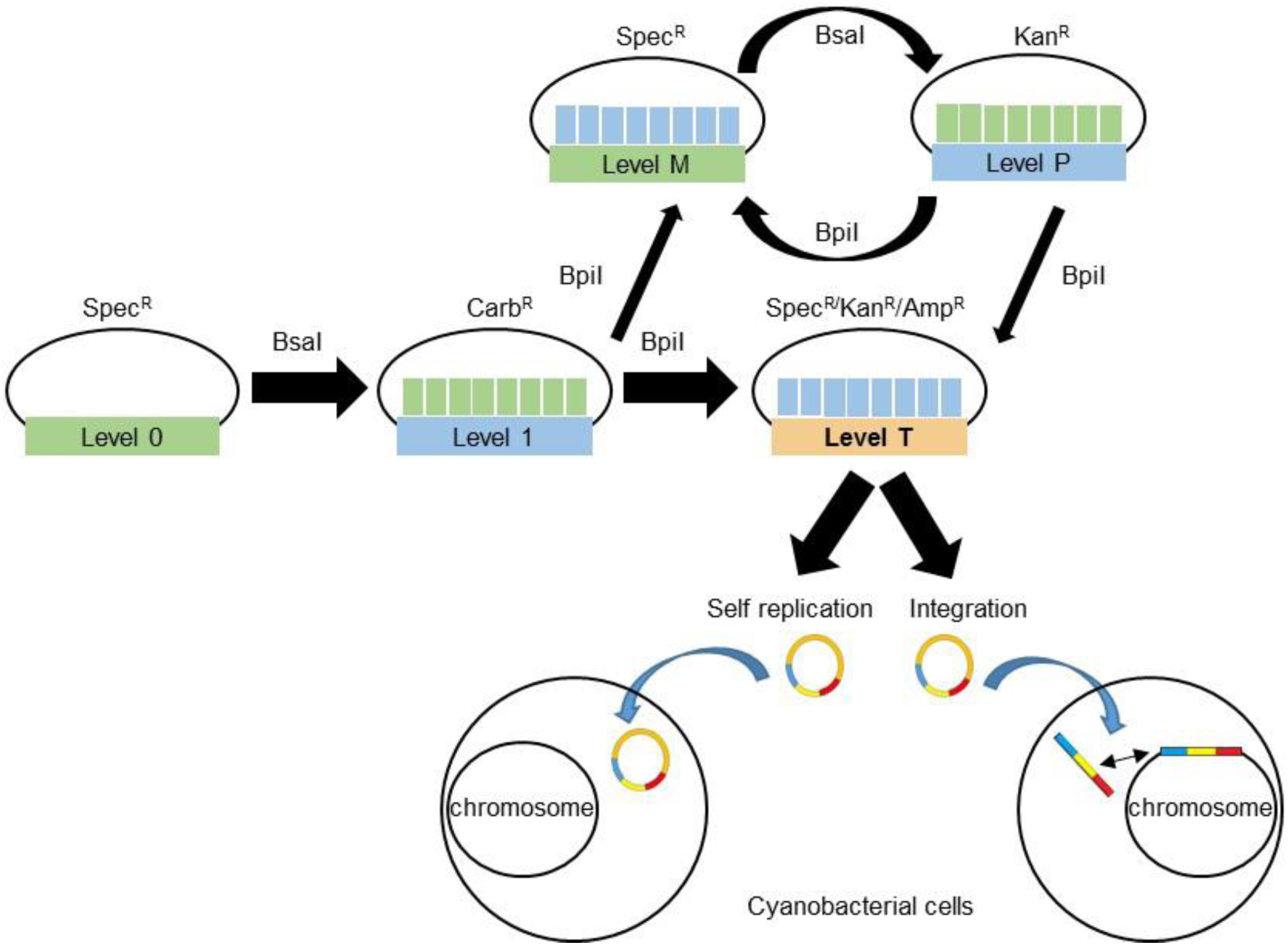
Extension of the Plant Golden Gate MoClo Assembly Standard for cyanobacterial transformation. Assembly relies one of two Type IIS restriction endonuclease enzymes (*Bsa*I or *Bpi*I). Domesticated level 0 parts are assembled into level 1 vectors. Up to seven level 1 modules can be assembled directly into a level T cyanobacterial transformation vector, which consist of two sub-types (either a replicative or an integrative vector). Alternatively, larger vectors with more modules can be built by assembling level 1 modules into level M, and then cycling assembly between level M and level P, and finally transferred from Level P to level T. Antibiotic selection markers are shown for each level. Level T vectors are supplied with internal antibiotic selection markers (shown), but additional selection markers could be included from level 1 modules as required. See **Supp. Info. 3** for full list and maps of level T acceptor vectors.

#### Level 0 parts for CRISPRi and srRNA

A nuclease deficient Cas9 gene sequence sourced from Addgene (www.addgene.org/44249/) was domesticated and assembled as a level 0 CDS part (Supp. Info. 1;2) (52). Five promoters of different strengths were truncated to the transcriptional start site (TSS) and cloned into a new level 0 acceptor vector with the unique overhangs GGAG and TAGC (Fig. 1). Two new level 0 parts with the unique overhangs GTTT and CGCT were generated for the sgRNA scaffold and srRNA HFQ handle (based on MicC) (43), respectively. Assembly of level 1 expression cassettes proceeded by combining appropriate level 0 parts with a PCR product for either a srRNA or sgRNA (Fig. 1).

### Cyanobacterial transformation and conjugation

Transformation with integrative level T vectors was performed as in Lea-Smith et al. (20). For transformation by electroporation, cultures were harvested during the exponential growth phase (OD_750_ of ~0.6) by centrifugation at 4,000 *g* for 10 min. The cell pellet was washed 3 times with 2 ml of sterile 1 mM HEPES buffer (pH 7.5), re-suspended in water with 3-5 μg of level T vector DNA and transferred into a 0.1 cm electroporation cuvette (Scientific Laboratory Suppliers). Re-suspended cells were electroporated using an Eppendorf 2510 electroporator (Eppendorf) set to 1200 V. Sterile BG-11 (1 ml) was immediately added to the electroporated cells. Following a 1 hr incubation at RT, the cells were plated on 1.5% (w/v) agar plates containing BG-11 with antibiotics at standard working concentrations to select for transformed colonies. The plates were sealed with parafilm and placed under 15 μmol photons m^−2^ s^−1^ light at 30°C for 1 day. The plates were then moved to 30 μmol photons m^−2^ s^−1^ light until colonies appeared. After 15-20 days, putative transformants were recovered and streaked onto new plates with appropriate antibiotics for further study.

Genetic modification by conjugation in *Synechocystis* was facilitated by an *E. coli* strain (HB101) carrying both mobilizer and helper vectors pRK2013 (ATCC^®^ 37159™) and pRL528 (www.addgene.org/58495/), respectively (53). For UTEX 2973, conjugation was facilitated by a HB101 strain carrying mobilizer and helper vectors pRK24 (www.addgene.org/51950/) and pRL258 (www.addgene.org/58495/), respectively. Cultures of HB101 and OneShot TOP10 *E. coli* strains carrying level T cargo vectors were grown for approximately 15 hr with appropriate antibiotics. Cyanobacterial strains were grown to an OD_750_ of ~1. All bacterial cultures were washed three times with either fresh LB medium for *E. coli* or BG11 for cyanobacteria prior to use. *Synechocystis* cultures (100 μl, OD_750_ of 0.5-0.8) were conjugated by combining appropriate HB101 and the cargo strains (100 μl each) and plating onto HATF 0.45 μm transfer membranes (Merck Millipore) placed on LB: BG11 (1: 19) agar plates. For UTEX 2973 conjugations, appropriate HB101 and the cargo strains (100 μl each) were initially combined and incubated at 30°C for 30 min, then mixed with UTEX 2973 cultures (100 μl, OD_750_ of 0.5-0.8) and incubated at 30°C for 2 hr, and then plated onto transfer membranes as above. *Synechocystis* and UTEX 2973 transconjugates were grown under culturing conditions outlined above. Following growth on non-selective media for 24 hr, the membranes were transferred to BG11 agar plates supplemented with appropriate antibiotics. Colonies were observed within a week for both strains.

### Cell counts, soluble protein and eYFP quantification

*Synechocystis* and UTEX 2973 strains were cultured for 48 hr as described above, counted using a haemocytometer and then harvested for soluble protein extraction. Cells were pelleted by centrifugation at 4,000 *g* for 15 min, re-suspended in lysis buffer [0.1 M potassium phosphate buffer (pH 7.0), 2 mM DTT and one Roche cOmplete EDTA-free protease inhibitor tablet per 10 ml (Roche Diagnostics)] and lysed with 0.5 mm glass beads (Thistle Scientific) in a TissueLyser II (Qiagen). The cell lysate was centrifuged at 18,000 g for 30 min and the supernatant assayed for soluble protein content using Pierce 660nm Protein Assay Reagent against BSA standards (Thermo Fisher Scientific). Extracts were subjected to sodium dodecyl sulfate–polyacrylamide gel electrophoresis (SDS-PAGE) in a 4–12% (w/v) polyacrylamide gel (Bolt^®^ Bis-Tris Plus Gel; Thermo Fisher Scientific) alongside a SeeBlue Plus2 Prestained protein ladder (Thermo), transferred to polyvinylidene fluoride (PVDF) membrane then probed with monoclonal anti-GFP serum (AbCAM) at 1: 1,000 dilution, followed by Li-Cor IRDye ^®^800CW goat anti-rabbit IgG (Li-Cor Inc.) at 1: 10,000 dilution, then viewed on an Li-Cor Odyssey CLx Imager. eYFP protein content was estimated by immunoblotting using densitometry using Li-Cor Lite Studio software v5.2. Relative eYFP protein abundance was estimated by densitometry using Li-Cor Lite Studio software v5.2.

### Confocal laser scanning microscopy

Cultures were imaged using Leica TCS SP8 confocal microscopy (Leica Microsystems) with a water immersion objective lens (HCX APO L 20×/0.50 W). Excitation/emission wavelengths were 514 nm/527-546 nm for eYFP and 514 nm/605−710 nm for chlorophyll autofluorescence.

## RESULTS AND DISCUSSION

### Construction of the CyanoGate system

The CyanoGate system integrates with the two-part Golden Gate MoClo Plant Tool Kit, which can be acquired from Addgene [standardised parts (Kit #1000000047) and backbone acceptor vectors (Kit # 1000000044), (www.addgene.org)] (38). The syntax for level 0 parts was adapted for prokaryotic cyanobacteria to address typical cloning requirements for cyanobacterial research (Fig. 1). New level 0 parts were assembled from a variety of sources (Supp. Info. 1). Level 1, M and P acceptor vectors were adopted from the MoClo Plant Tool Kit, which facilitates assembly of level 0 parts in a level 1 vector, and subsequently up to seven level 1 modules in level M. Level M assemblies can be combined further into level P and cycled back into level M to produce larger multi-module vectors if required (Supp. Info. 4). Vectors >50 kb in size assembled by MoClo have been reported (54). Modules from level 1 or level P can be assembled in new level T vectors designed for cyanobacterial transformation (Fig. 2). We found that both UTEX 2973 and *Synechocystis* produced recombinants following electroporation or conjugation methods with level T vectors. For the majority of the work outlined below, we relied on the conjugation approach.

### Integration - generating marked and unmarked knockout mutants

A common method for engineering stable genomic knock out and knock in mutants in several cyanobacteria relies on homologous recombination via integrative (suicide) vectors using a two-step marked-unmarked strategy (20) (Supp. Info. 5). Saar et al. (13) recently used this approach to introduce up to five genomic alterations into a single *Synechocystis* strain. Firstly, marked mutants are generated with an integrative vector carrying two sequences (approximately 1 kb each) identical to the regions of the cyanobacterial chromosome flanking the deletion/insertion site. Two gene cassettes are inserted between these flanking sequences: a levansucrase expression cassette (*sac*B) that confers sensitivity to transgenic colonies grown on sucrose and an antibiotic resistance cassette (Ab*^R^*) of choice. Secondly, unmarked mutants (carrying no selection markers) are generated from fully segregated marked lines using a separate integrative vector carrying only the flanking sequences and selection on plates containing sucrose.

We adapted this approach for the CyanoGate system (Fig. 1). To generate level 1 vectors for making knock out mutants, sequences flanking the upstream (UP FLANK) and downstream (DOWN FLANK) site of recombination were ligated into the plant MoClo Prom+5U (with overhangs GGAG-AATG), and 3U+Ter (GCTT-CGCT) positions, respectively, to generate new level 0 parts (Fig. 1B). In addition, full expression cassettes were made for sucrose selection (*sac*B) and antibiotic resistance (Ab*^R^*Spec, Ab*^R^*Kan and Ab*^R^*Ery) in level 0 that ligate into positions SP (AATG-AGGT) and CDS2 (stop) (AGGT-GCTT), respectively. Marked level 1 modules can be assembled using UP FLANK, DOWN FLANK, sacB and the required Ab*^R^* level 0 part. For generating the corresponding unmarked level 1 module, a short 59 bp linker (UNMARK LINKER) can be ligated into the CDS1ns (AATG-GCTT) position for assembly with an UP FLANK and DOWN FLANK (Fig. 1D). Unmarked and marked level 1 modules can then be assembled into level T integrative vectors, with the potential capacity to include multiple knockout modules in a single level T vector.

To validate our approach, we constructed the level 0 flanking vectors pC0.024 and pC0.025 and assembled two level T integrative vectors using pUC19-T (*cpcBA*-M and *cpcBA*-UM, with and without the *sac*B and Ab*^R^* cassettes, respectively) to remove the *cpcBA* promoter and operon in *Synechocystis* and generate an “Olive” mutant unable to produce the phycobiliprotein C-phycocyanin (55,56) (Fig. 3A, B; Supp. Info. 1). Following transformation with *cpcBA*-M, we successfully generated a marked Δ*cpcBA* mutant carrying the sacB and the Ab*^R^*Kan cassettes after selective segregation (*ca*. 3 months) (Fig. 3C). The unmarked Δ*cpcBA* mutant was then isolated following transformation of the marked Δ*cpcBA* mutant with *cpcBA*-UM and selection on sucrose (*ca*. 2 weeks).

**Figure 3.**
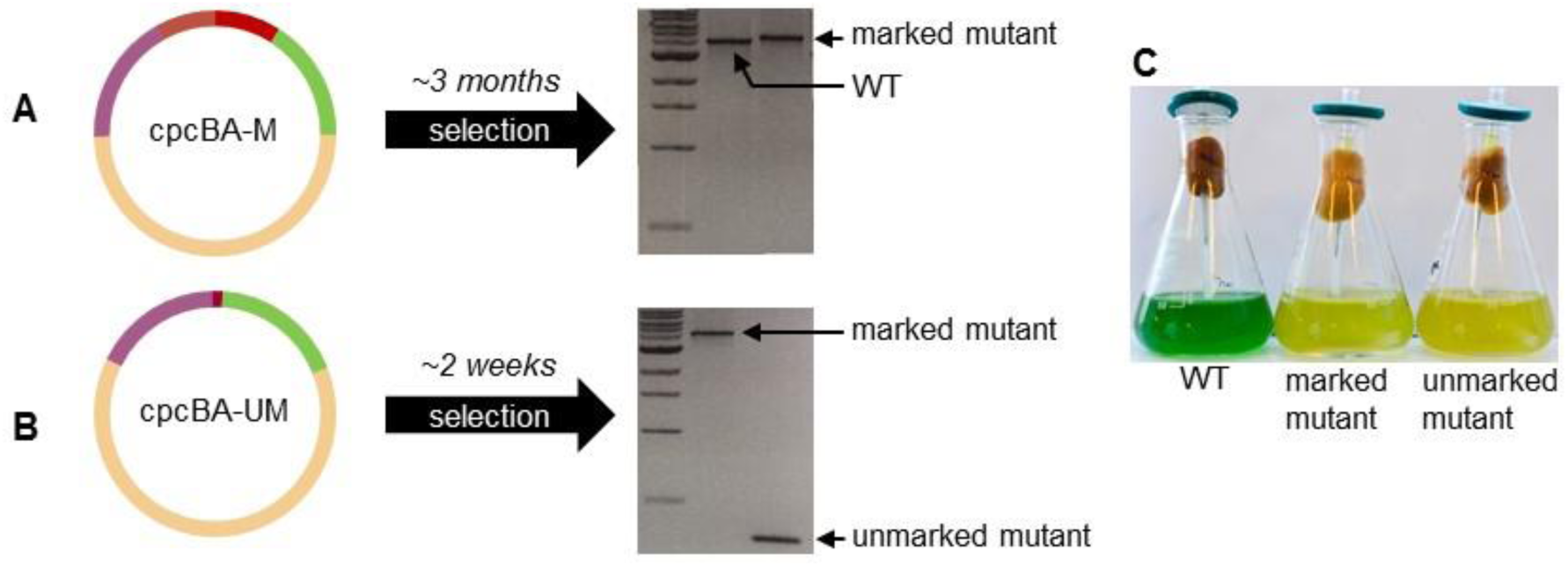
Generating knock out mutants in cyanobacteria. (A) Assembled level T vector *cpcBA*-M (see Fig. 1C) targeting the *cpcBA* promoter and operon (3,563 bp) to generate a marked Δ*cpcBA* “Olive” mutant in *Synechocystis* sp. PCC 6803. Following transformation and segregation on Kanamycin (ca. 3 months), a segregated marked mutant was isolated (WT band is 3,925 bp, marked mutant band is 5,503 bp, 1kb DNA ladder (NEB) is shown). (B) Assembled level T vector *cpcBA*-UM (see Fig. 1D) for generating an unmarked Δ*cpcBA* mutant. Following transformation and segregation on sucrose (*ca*. 2 weeks), an unmarked mutant was isolated (unmarked band is 425 bp). (C) Liquid cultures of WT, marked and unmarked Olive mutants.

### Generating knock in mutants

Flexibility in designing level 1 insertion cassettes is needed when making knock in mutants. Thus, for knock in mutants the upstream and downstream sequences flanking the insertion site, and any required expression or marker cassettes, are first assembled into separate level 1 modules from UP FLANK and DOWN FLANK level 0 parts (Fig. 1E, F). Seven level 1 modules can be assembled directly into Level T (Fig. 2), thus with a single pair of flanking sequences up to five level 1 expression cassettes could be included in a Level T vector.

Linker parts (20 bp) UP FLANK LINKER and DOWN FLANK LINKER were generated to allow assembly of level 0 UP FLANK and DOWN FLANK parts into separate level 1 acceptor vectors. Similarly, level 0 linker parts were generated for *sac*B and Ab*^R^* (Fig. 1H, I). Level 1 vectors at different positions can then be assembled in level T (or M) containing one or more expression cassettes, an Ab*^R^* of choice, or both *sac*B and Ab*^R^* (Fig. 2).

Using this approach, CyanoGate can facilitate the generation of knock in mutants using a variety of strategies. For example, if retention of a resistance marker is not an experimental requirement (e.g. 57), only a single antibiotic resistance cassette needs to be included in level T. Alternatively, a two-step marked-unmarked strategy could be followed, as for generating knockout mutants.

While knock out strategies can target particular loci, knock in approaches often rely on recombination at previously identified neutral sites within the genome of interest. We have assembled a suite of neutral site flanking regions to target neutral sites in *Synechocystis* (56, 58), *Synechococcus* sp. PCC 7002 (PCC 7002 hereafter) (59), and neutral sites common to UTEX 2973, PCC 7942 and *Synechocystis* sp. PCC 6301 (60-62). (Supp. Info. 1).

To validate our system, we generated a level T vector carrying the flanking regions for the *cpcBA* operon and an eYFP expression cassette (*cpcBA*-eYFP) (Fig. 4A, B; Supp. Info. 1). We successfully transformed this vector into our marked “Olive” *Synechocystis* mutant, to generate a stable olive mutant with constitutive expression of eYFP (Olive-eYFP) (Fig. 4C).

**Figure 4.**
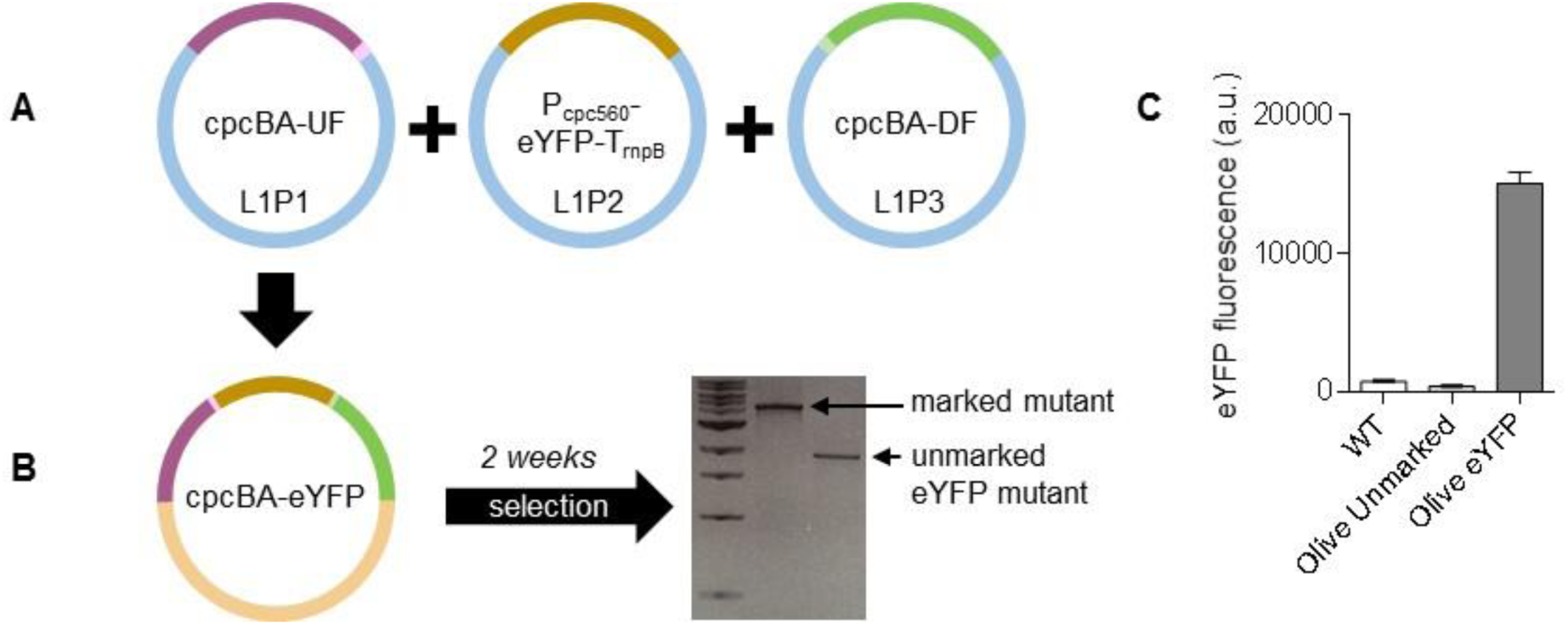
Generating knock in mutants in cyanobacteria. (A) Assembly of level 1 modules *cpcBA-*UF (see Fig. 1E) in the level 1, position 1 acceptor (L1P1), P*_cpc560_-eYFP-*T*_rnbP_* (see Fig. 1G) in L1P2 and *cpcBA*-DF (see Fig. 1F) in L1P3. (B) Transfer of level 1 assemblies to level T vector *cpcBA*-eYFP for generating an unmarked Δ*cpcBA* mutant carrying an eYFP expression cassette. Following transformation and segregation on sucrose (<u>ca</u>. 3 weeks), an unmarked eYFP mutant was isolated (1,771 bp). (C) Fluorescence values are the means ± SE of four biological replicates.

### Expression comparison for promoter parts in *Synechocystis* and UTEX 2973

We constructed level 0 parts for a wide selection of synthetic promoters and promoters native to *Synechocystis*. Promoters were assembled as expression cassettes driving eYFP in replicative level T vector pPMQAK1-T to test for differences in expression when conjugated into *Synechocystis* or UTEX 2973. We first compared the growth rates of *Synechocystis*, UTEX and PCC 7942 [a close relative of UTEX 2973 (46)] under a variety of different culturing conditions (Supp. Info. 6). We found that growth rates were comparable between *Synechocystis* and PCC 7942 at temperatures below 40°C regardless of light levels and supplementation with CO_2_. In contrast, UTEX 2973 grew poorly under those conditions. UTEX 2973 only showed an enhanced growth rate at 45°C under the highest light tested (500 μmol photons m^−2^ s^−1^) with CO_2_, while all three strains failed to grow at 50°C. These results confirm that the fast-growth phenotype reported for UTEX 2973 requires specific conditions as reported by Ungerer et al. (47). We proceeded with CyanoGate part characterisations and comparisons under the best conditions achievable for *Synechocystis* and UTEX 2973 (see Materials and Methods) (Supp. Info. 7A).

#### Promoters native to *Synechocystis*

We assembled several previously reported promotors from *Synechocystis* in the CyanoGate kit. These include six inducible/repressible promoters (P*_nrsB_*, P*_coaT_*, *P_nirA_, P_petE_*, P*_isiAB_*, and P*_arsB_*), which were placed in front a strong ribosomal binding site (RBS^*^) (Supp. Info. 1; 2) (16, 27). P*_nrsB_* and P*_coaT_* drive the expression of nickel and cobalt ion efflux pumps and are induced by Ni^2+^, and Co^2+^ or Zn^2+^, respectively (27,63-65). P_nirA_, from the nitrate assimilation operon, is induced by the presence of NO_3_− and/or NO_2_^−^ (52,66). P_petE_ drives the expression of plastocyanin and is induced by Cu^2+^, which has previously been used for the expression of heterologous genes (24,65). The promoter of the *isiAB* operon (P*_isiAB_*) is repressed by Fe^3+^ and activated when the cell is under iron stress (67). P*_arsB_* drives the expression of a putative arsenite and antimonite carrier and is activated by AsO_2_^−^ (64).

We also cloned the *rnpB* promoter, P*_rnpB_*, from the Ribonuclease P gene (18), a long version of the *psbA2* promoter, P*_psbA2L_*, from the Photosystem II protein D1 gene (27,68) and the promoter of the C-phycocyanin operon, P*_cpc560_* (also known as P*_cpcB_* and P*_cpcBA_*) (69). P*_rnpB_* and P*_psbA2L_* were placed in front of RBS^*^ (16) (Fig. 5A). For P*_cpc560_* we assembled four variants. Firstly, P*_cpc560+A_* consisted of the promoter and the 4 bp MoClo overhang AATG. Secondly, P*_cpc560_* was truncated by one bp (A), so that that the start codon was aligned with the native P*_cpc560_* RBS spacer region length. Zhou et al. (69) identified 14 predicted transcription factor binding sites (TFBSs) in the upstream region of P*_cpc560_* (-556 to -381 bp) and removal of this region resulted in a significant loss of promoter activity. However, alignment of the reported TFBSs showed their locations are in the downstream region of the promoter (-180 to -5 bp). We identified 11 additional predicted TFBSs using Virtual Footprint (70) in the upstream region and hypothesised that the promoter activity may be modified by duplicating either of these regions. So thirdly, we generated P*_cpc560_Dx2_*containing a duplicated downstream TFBS region. For P*_cpc560_Dx2_*, only the region between -31 to -180 bp was duplicated to avoid repeating the Shine-Dalgarno (SD) sequence. Fourthly, we duplicated the upstream region to generate P*_cpc560_Ux2_*. We then assembled P*_rnpB_*, P*_psbA2L_* and the four P*_cpc560_* variants with eYFP and the *rnpB* terminator (T*_rnpB_*) into a Level 1 expression cassette, and subsequently into a level T replicative vector (pPMQAK1-T) for expression analysis (Supp. Info. 8).

**Figure 5.**
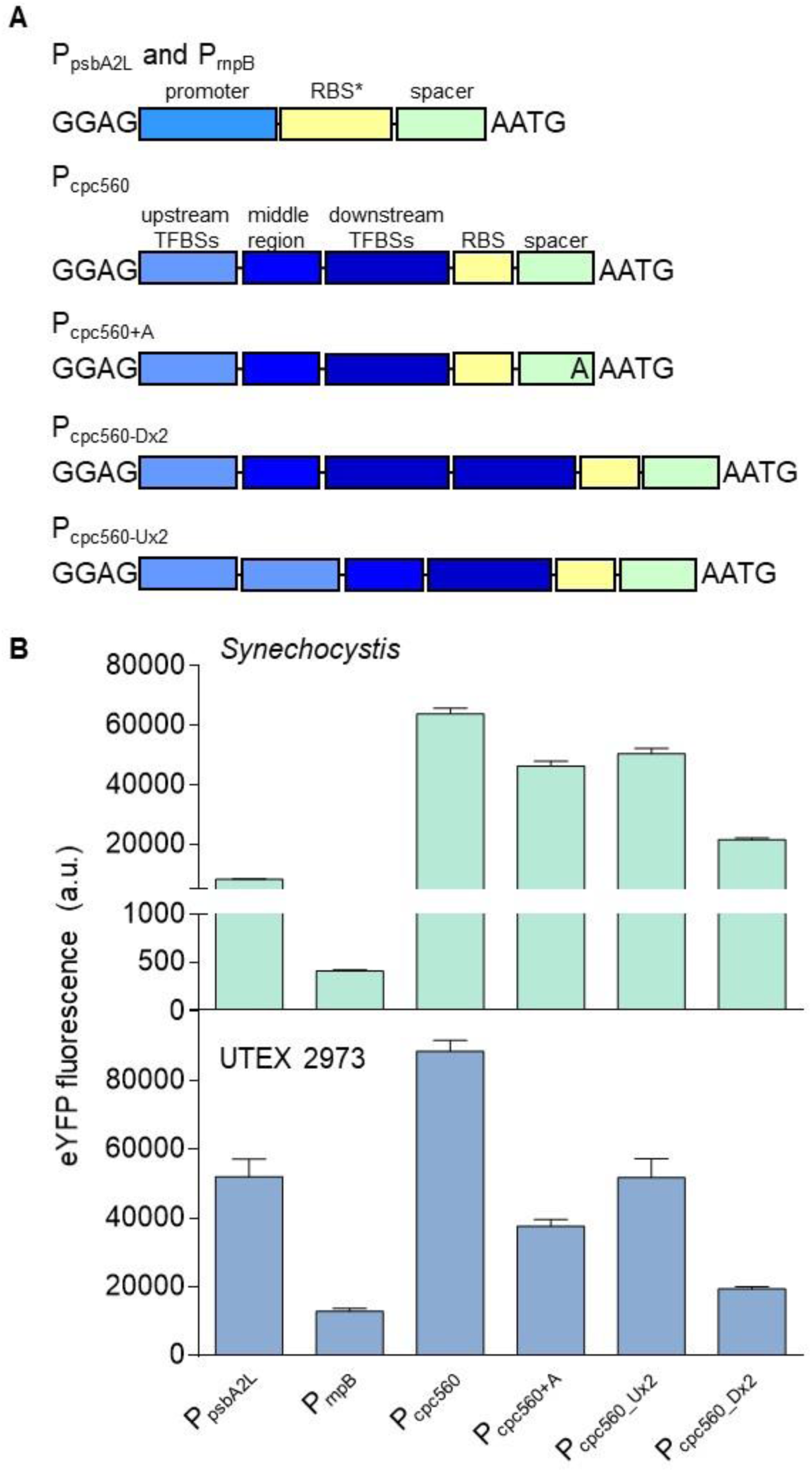
Expression levels of cyanobacterial promoters in *Synechocystis* and UTEX 2973. (A) Structure of the cyanobacterial promoters adapted for the CyanoGate kit. Regions of P*_cpc560_* shown are the upstream transcription factor binding sites (TFBSs) (-556 to -381 bp), middle region (-380 to -181 bp), and the downstream TFBSs, ribosome binding site (RBS) and spacer (-180 to -5 bp) (B) Expression levels of eYFP driven by promoters in *Synechocystis* and UTEX 2973. Values are the means ± SE from at least four biological replicates.

In *Synechocystis* the highest expressing promoter was P*_cpc560_* (Fig. 5B), which indicated that maintaining the native RBS spacer region for P*_cpc560_* is important for maximising expression. Neither P*_cpc560_Dx2_* nor P*_cpc560_Ux2_* had higher expression levels compared to P*_cpc560_*. P*_cpc560_Dx2_* was strongly decreased compared to P_cpc560_, suggesting that promoter function is sensitive to modification of the downstream region and this region could be a useful target for modulating P*_cpc560_* efficacy. Previous work in *Synechocystis* has suggested that modification of the middle region of P*_cpc560_* (-380 to -181 bp) may also affect function (56). P*_psbA2L_* produced lower expression levels than any variant of P*_cpc560_* in *Synechocystis*, while P*_rnpB_* produced the lowest expression levels. The observed differences in expression levels are consistent with those in other studies with *Synechocystis* (24, 27, 33).

In UTEX 2973, the trend in expression patterns was similar to that in *Synechocystis* (Fig. 5B). However, the overall expression levels of eYFP measured in UTEX 2973 were significantly higher than in *Synechocystis*. P*_cpc560_* was increased by 30%, while P*_rnpB_* showed a 20-fold increase in expression relative to *Synechocystis*. The relative expression strength of P*_psbA2L_* was also higher than in *Synechocystis*, and second only to P*_cpc560_* in UTEX 2973. As promoters derived from PpsbA are responsive to increasing light levels (27), the increased levels of expression for P_psbA2L_ may be associated with the higher light intensities used for growing UTEX 2973 compared to *Synechocystis*. Background fluorescence levels were similar between UTEX 2973 and *Synechocystis* conjugated with an empty pPMQAK1 -T vector, which suggested that the higher fluorescence values in UTEX 2973 were a direct result of increased levels of eYFP protein.

#### Heterologous and synthetic promoters

A suite of twenty constitutive synthetic promoters was assembled in level 0 based on the modified BioBricks BBa_J23119 library of promoters (26), and the synthetic P*_trc10_*, P*_tic10_* and P*_tac10_* promoters (18, 25) (Supp. Info. 1; 2). We retained the broad-range BBa_B0034 RBS (AAAGAGGAGAAA) and *lac* operator (*lacO*) from Huang et al. (18), for future *lacI-based* repression experiments (*lacI* and the P_lacIQ_ promoter are included in the CyanoGate kit) (71). We cloned eight new variants (J23119MH_V01-8) with mutations in the canonical BBa_J23119 promoter sequence (Fig. 6A). Additionally, we included the L-arabinose-inducible promoter from *E. coli* (P*_BAD_*) (72).

**Figure 6.**
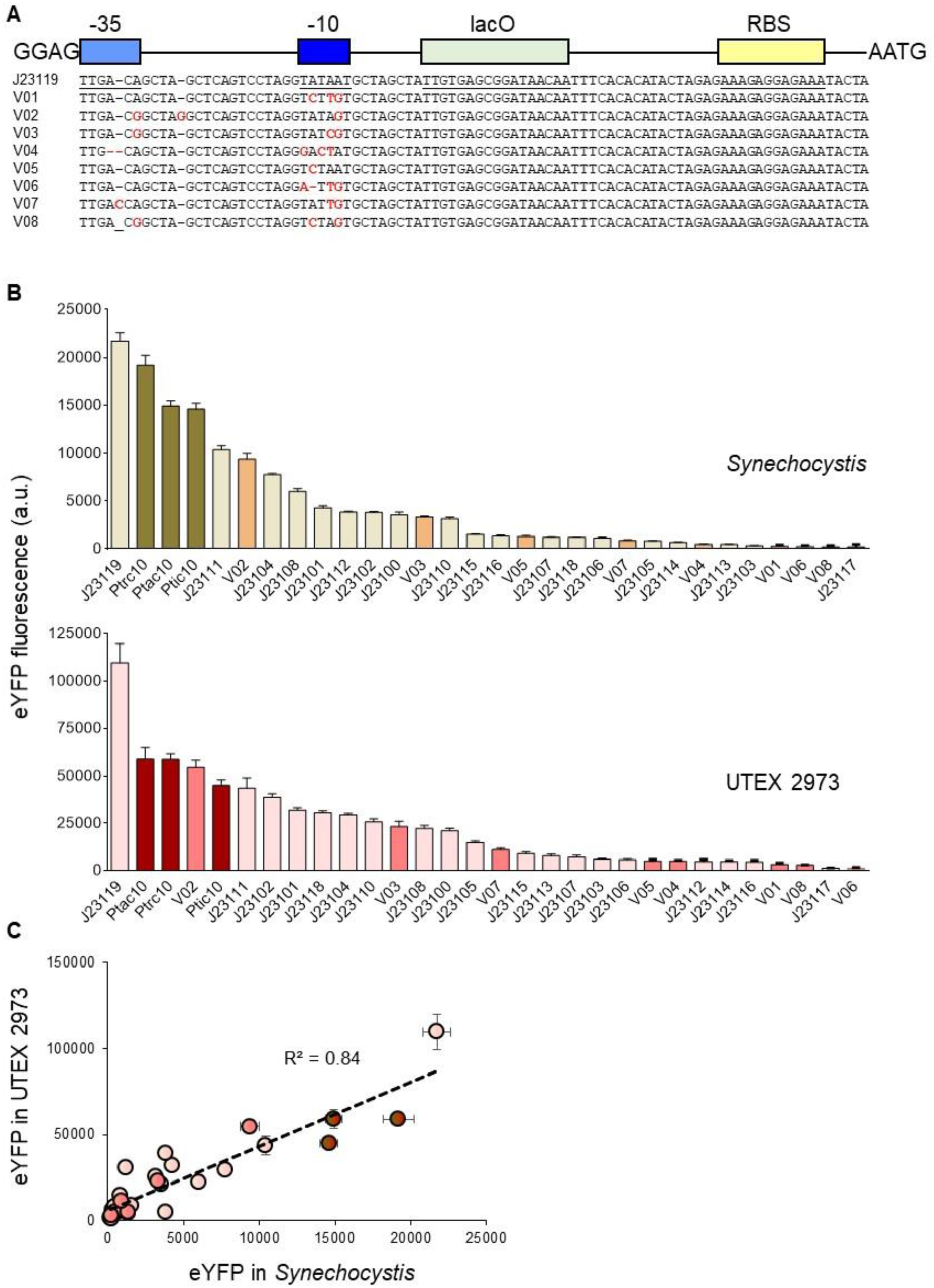
Expression levels of heterologous and synthetic promoters in *Synechocystis* and UTEX 2973. (A) Structure and alignment of eight new synthetic promoters derived from the BioBricks BBa_J23119 library and *P_trc10_* promoter design (18). (B) Expression levels of eYFP driven by promoters in *Synechocystis* and UTEX 2973. (C) Correlation analysis of expression levels of synthetic promoters tested in *Synechocystis* and UTEX 2973. The coefficient of determination (R^2^) is shown for the J23119 library (red), new synthetic promoters (pink) and *trc* variants (dark red). Values are the means ± SE from at least four biological replicates. See **Supp. Info. 7** for more info.

We encountered an unexpected challenge with random internal deletions in the -35 and -10 regions of some promoters of the BBa_J23119 library and *trc* promoter variants when cloning them into level 0 acceptors. Similar issues were reported previously for the *E. coli* EcoFlex kit (36) that may relate to the functionality of the promoters and the host vector copy number in *E. coli*, which consequently resulted in cell toxicity and selection for mutated promoter variants. To resolve this issue, we generated a low copy level 0 promoter acceptor vector compatible with CyanoGate (pSB4K5 acceptor) for cloning recalcitrant promoters (Supp Info. 1; 2). Subsequent assemblies in level 1 and T showed no indication of further mutation.

We then tested the expression levels of eYFP driven by the synthetic promoters in *Synechocystis* and UTEX 2973 following assembly in pPMQAK1-T (Fig. 6B; Supp. Info. 8). The synthetic promoters showed a 120-fold dynamic range in both cyanobacterial strains. Furthermore, a similar trend in promoter expression strength was observed (R^2^ = 0.84) (Fig. 6C). However, eYFP expression levels were on average eight-fold higher in UTEX 2973 compared to *Synechocystis*. In *Synechocystis*, the highest expression levels were observed for J23119 and P*_trc10_*, but these were still approximately 50% lower than values for the native P*_cpc560_* promoters (Fig. 5B). The expression trends for the BBa_J23119 library were consistent with the subset reported by Camsund et al. (24) in *Synechocystis*, while the observed differences between P*_trc10_* and P*_cpc560_* were similar to those reported by Lui et al. (33).

In contrast, the expression levels in UTEX 2973 for J23119 were approximately 50% higher than P*_cpc560_*. Several synthetic promoters showed expression levels in a similar range to those for the native P*_cpc560_* promoter variants, including P*_trc10_*, J23111 and the J23119 variant V02. V02 is identical to J23111 except for an additional ‘G’ between the -35 and -10 motifs, suggesting that small changes in the length of this spacer region may not be critical for promoter strength (similar expression levels were also observed for these two promoters in *Synechocystis*). In contrast, a single bp difference between J23111 and J23106 in the -35 motif resulted in an eight- and ten-fold reduction in expression in *Synechocystis* and UTEX 2973, respectively. The results for UTEX 2973 were unexpected, and to our knowledge no studies to date have directly compared these promoters in this strain. Recent work has examined the expression of β-galactosidase using promoters such as P*_cpc560_* and P*_trc_* in UTEX 2973 (32). Li et al. (32) highlighted that different growth environments (e.g. light levels) can have significant effects on protein expression. Changes in culture density can also affect promoter activity (73), so we tracked eYFP expression levels over time for three days for *Synechocystis* and UTEX 2973. Although expression levels for each promoter fluctuated over time, with peak expression levels at 24 hr and 48 hr in UTEX 2973 and *Synechocystis*, respectively, the overall expression trends were generally consistent for the two strains (Supp. Info. 7B).

### Protein expression levels in *Synechocystis* and UTEX 2973

To investigate further the increased levels of eYFP expression observed in UTEX 2973 compared to *Synechocystis*, we examined cell morphology, protein content and eYFP protein abundances in expression lines for each strain. Confocal image analysis confirmed the coccoid and rod shapes of *Synechocystis* and UTEX 2973, respectively, and the differences in cell size (46, 74) (Fig. 7A). Immunoblot analyses of eYFP from protein extracts of four eYFP expressing strains correlated well with previous flow cytometry measurements (Fig 5; 6). eYFP driven by the J23119 promoter in UTEX 2973 produced the highest levels of eYFP protein (Fig. 7B, C). Although the density of cells in culture was two-fold higher in *Synechocystis* compared to UTEX 2973 (Fig. 7D), the protein content per cell was six-fold lower (Fig. 7E). We then estimated the average cell volumes for *Synechocystis* and UTEX 2973 at 3.8 ± 0.12 μm^3^ and 6.7 ± 0.25 μm^3^ (n = 30 each), respectively, based on measurements from confocal microscopy images. Based on those estimates, we calculated that the density of soluble protein per cell was four-fold higher in UTEX 2973 compared to *Synechocystis* (Fig. 7F). Thus, we hypothesised that the enhanced levels of eYFP observed in UTEX 2973 were a result of the expression system harnessing a larger available amino acid pool. Mueller et al. (75) have reported that UTEX 2973 has an increased investment in amino acid content compared to PCC 7942, which may be linked to higher rates of translation in UTEX 2973. Therefore, UTEX 2973 continues to show promise as a bioplatform for generating heterologous protein products, although future work should study production rates under conditions optimal for faster-growth (47).

**Figure 7.**
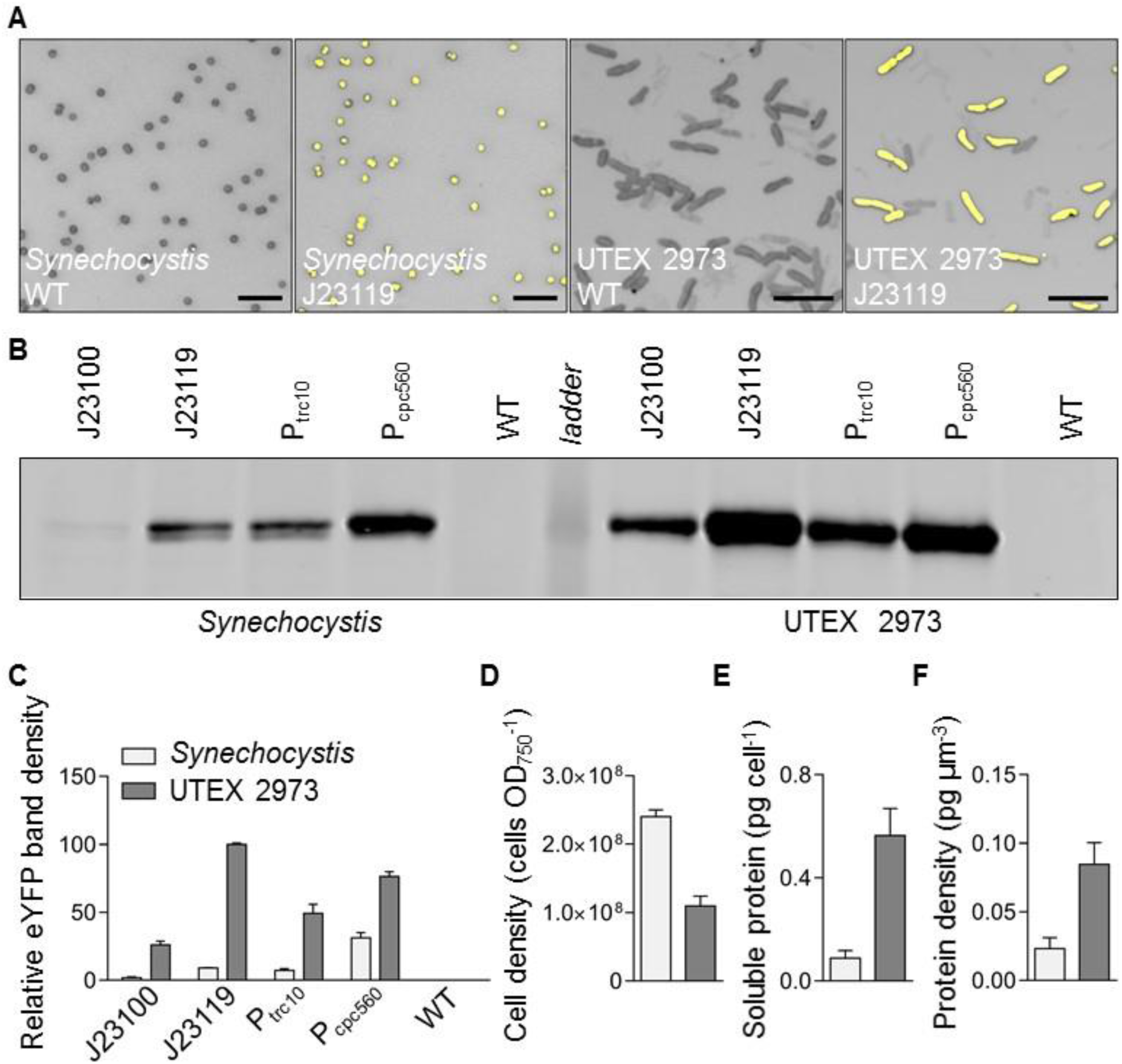
Protein expression levels in *Synechocystis* and UTEX 2973 cells. (A) Confocal images of WT strains and mutants expressing eYFP driven by the J23119 promoter (bar = 10 μm). (B) Representative immunoblot of protein extracts (3 μg protein) from mutants with different promoter expression cassettes (as in **Fig. 6**) probed with an antibody against eYFP. The protein ladder band corresponds to 30 kDa. (E) Relative eYFP protein abundance relative to UTEX 2973 mutants carrying the J23119 expression cassette. (C-E) Cell density, protein content per cell and protein density per estimated cell volume for *Synechocystis* and UTEX 2973. Values are the means ± SE of four biological replicates.

### The RK2 origin of replication is functional in *Synechocystis*

Synthetic biology tools (e.g. gene expression circuits, CRISPR/Cas-based systems) are often distributed between multiple plasmid vectors at different copy numbers in order to synthesise each component at the required concentration (76). The large RSF1010 vector is able to replicate in a broad range of microbes including gram-negative bacteria such as *E. coli* and several cyanobacterial species. However, for 25 years it has remained the only non-native vector reported to be able to self-replicate in cyanobacteria (17). Recently, two small plasmids native to *Synechocystis*, pCA2.4 and pCB2.4, have been engineered for gene expression (33, 73, 77). The pANS plasmid (native to PCC 7942) has also been adapted as a replicative vector, but so far it has been only shown to function in PCC 7942 and *Anabaena* PCC 7120 (78). To expand the replication origins available for cyanobacterial research further we tested the capacity for vectors from the SEVA library to replicate in *Synechocystis* (50).

We acquired three vectors driven by three different replication origins [pSEVA421 (RK2), pSEVA431 (pBBR1) and pSEVA442 (pRO1600/ColE1)] and carrying a spectinomycin antibiotic resistance marker. These vectors were domesticated and modified as level T acceptor vectors, assembled and then transformed into *Synechocystis* by electroporation or conjugation. Only *Synechocystis* strains conjugated with vectors carrying RK2 (pSEVA421-T) grew on Spectinomycin-containing plates (Supp. Info. 1; 2). We then assembled two level T vectors with an eYFP expression cassette (P*_cpc560_*-eYFP-T*_rnpB_*) to produce pPMQAK1-T-eYFP and pSEVA421-T-eYFP, which were conjugated into *Synechocystis* (Fig. 8; Supp. Info. 8). Both pPMQAK1-T and pSEVA421-T transconjugates grew at similar rates in 50 μg ml^−1^ Kanamycin and 5 μg ml^−1^ Spectinomycin, respectively. However, eYFP levels were 8-fold lower in pSEVA421-T-eYFP, suggesting that RK2 has a reduced copy number relative to RSF1010 in *Synechocystis*. This is consistent with reported lower copy numbers per cell for RK2 (4-7 copies) compared to RSF1010 (10-12 copies) in *E. coli* (79, 80).

**Figure 8.**
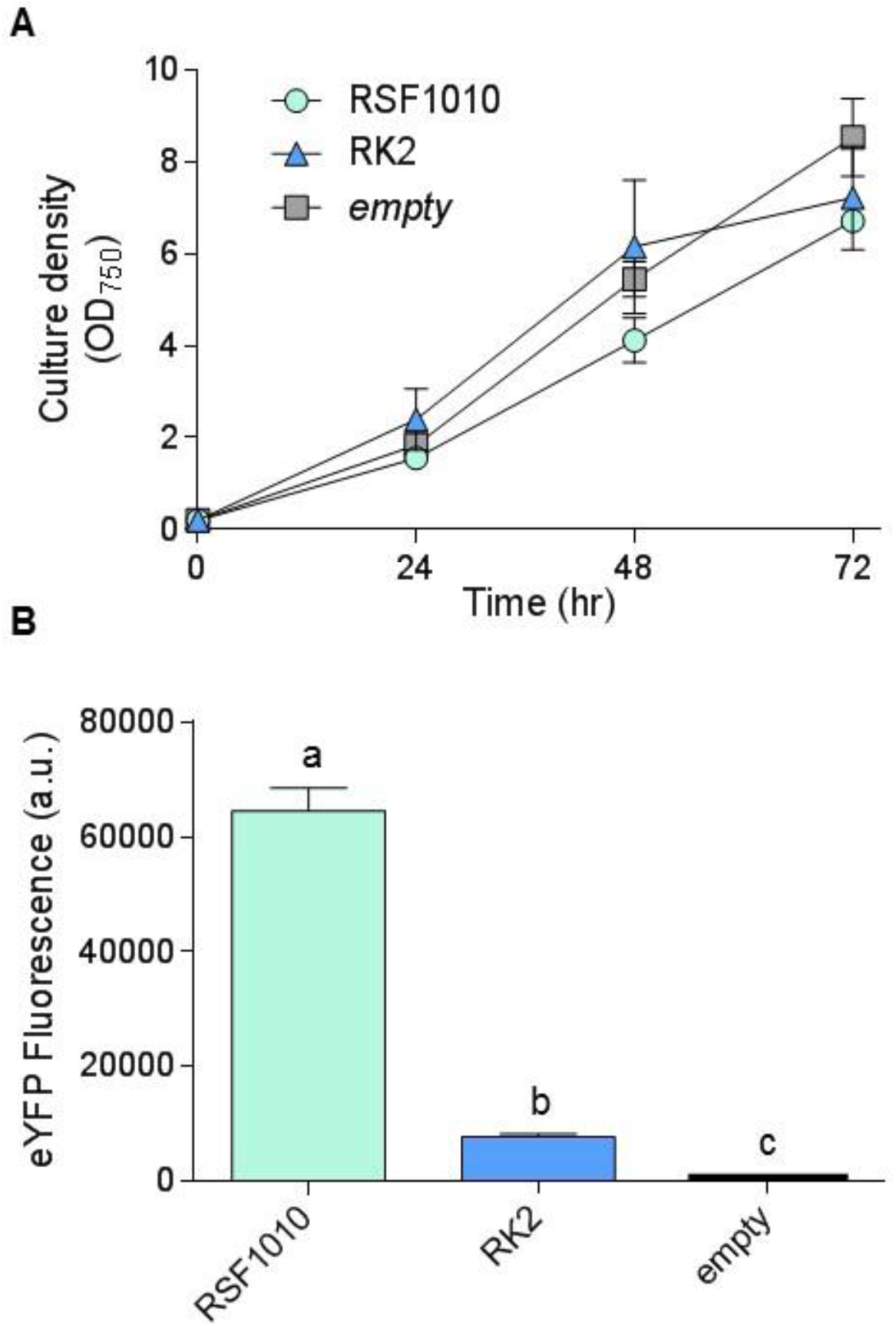
Growth and expression levels of eYFP with the RK2 replicative origin in *Synechocystis*. (A) Growth of strains carrying RK2 (vector pSEVA421-T-eYFP), RSF1010 (pPMQAK1-T-eYFP) or an empty pPMQAK1-T grown in appropriate antibiotics. Growth was measured as OD_750_ under a constant illumination of 100 μmol photons m^−2^s^−1^ at 30 °C. (B) Expression levels of eYFP after 48 hr of growth. Letters indicating significant difference (P < 0.05) are shown, as determined by ANOVA followed by Tukey’s HSD tests. Values are the means ± SE of four biological replicates.

### Gene repression systems

CRISPR (clustered regularly interspaced short palindromic) interference (CRISPRi) is a relatively new but well characterised tool for modulating genes expression at the transcription stage in a sequence-specific manner (81, 82). CRISPRi typically uses a nuclease deficient Cas9 from *Streptococcus pyogenes* (dCas9) and has been demonstrated to work in several cyanobacterial species, including *Synechocystis* (44), PCC 7002 (Gordon et al., 2016); PCC 7942 (83) and *Anabaena* sp. PCC 7120 (84). A second approach for gene repression uses rationally designed small regulatory RNAs (srRNAs) to regulate gene expression at the translation stage (43, 84). The synthetic srRNA is attached to a scaffold to recruit the Hfq protein, an RNA chaperone that is conserved in a wide-range of bacteria and cyanobacteria, which facilitates the hybridization of srRNA and target mRNA, and directs mRNA for degradation. The role of cyanobacterial Hfq in interacting with synthetic srRNAs is still unclear (85). However, regulatory ability can be improved by introducing Hfq from *E. coli* into *Synechocystis* (86). Both CRISPRi- and srRNA-based systems have potential advantages as they can be used to repress multiple genes simultaneously.

To validate the CRISPRi system, we assembled an expression cassette for dCas9 (P*_cpc560_*-dCas9-T*_rnpB_*) on the Level 1 position 1 vector pICH47732, and four different sgRNA expression cassettes (P*_trc10_*__TSS_-sgRNA-sgRNA scaffold) targeting eYFP on the Level 1 position 2 vector pICH47742 (38) (Fig. 9B, C; Supp. Info. 8). For assembly of CRISPRi sgRNA expression cassettes in level 1, we targeted four 18-22 bp regions of the eYFP non-template strand with an adjacent 3’ protospacer adjacent motif (PAM) of 5’-NGG-3’, as required by *S. pyogenes* dCas9. The sgRNA sequences contained no off-target sites in the *Synechocystis* genome [confirmed by CasOT (87)]. The sgRNAs were made by PCR using two complementary primers carrying the required overhangs and BsaI sites, and were assembled with P*_trc10_*_TSS promoter (pC0.203) and the sgRNA scaffold (pC0.122) (Fig. 9B). Level T vectors were assembled carrying dCas9 and a single sgRNA, or just the sgRNA alone. We subsequently conjugated the Olive-eYFP mutant and tracked eYFP expression.

**Figure 9.**
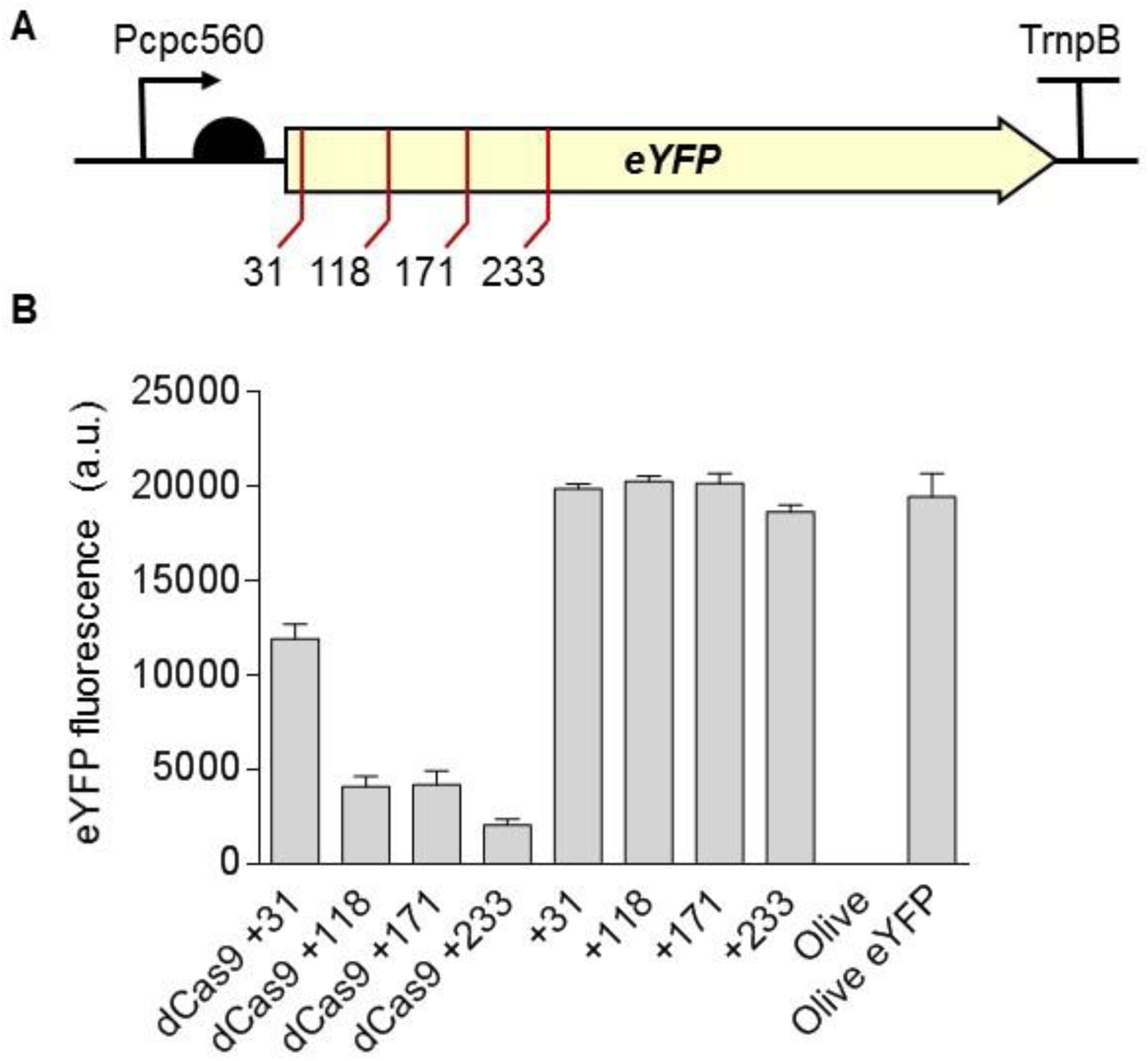
Gene regulation system using CRISPRi in *Synechocystis*. (A) Four target regions were chosen as sgRNA protospacers to repress eYFP expression in Olive-eYFP (**Fig. 4**): ‘CCAGGATGGGCACCACCC’ (+31), ‘ACTTCAGGGTCAGCTTGCCGT’ (+118), ‘AGGTGGTCACGAGGGTGGGCCA’ (+171) and ‘AGAAGTCGTGCTGCTTCATG’ (+233). (B) eYFP fluorescence of Olive-eYFP expressing constructs carrying sgRNAs with and without dCas9. Untransformed Olive-eYFP and the Olive mutant were used as controls. Values are the means ± SE of four biological replicates.

Transconjugates carrying only the sgRNA showed no reduction in eYFP level compared to non-transconjugated Olive-eYFP (Fig. 9D). However, all strains carrying dCas9 and a sgRNA showed a decrease in eYFP that ranged from 40-90% depending on the sgRNA used. These reductions are similar to those observed previously in PCC 7002 and in *Synechocystis* (44, 88) and demonstrated that CRISPRi system is functional in the CyanoGate kit.

## CONCLUSION

The CyanoGate kit was designed to increase the availability of well characterised libraries and standardised modular parts in cyanobacteria (45). We aimed to simplify and accelerate modular cloning methods in cyanobacterial research and allow integration with the growing number of labs that rely on the established common plant and algal syntax for multi-part DNA assembly (40, 42). Here, we have demonstrated the functionality of CyanoGate in sufficient detail to show that it is straightforward to adopt and functionally robust across different cyanobacterial species. CyanoGate could likely be utilised also in non-cyanobacterial microbes amenable to transformation (e.g. *Rhodopseudomonas* spp.) and adapted for use in subcellular eukaryotic compartments of prokaryote origin (e.g. chloroplasts) (89-91). In addition to the parts discussed, we have also assembled a suite of 21 terminators (Supp. Info. 1). To increase the accessibility and usability of the CyanoGate, we have included the vector maps for all parts and new acceptors (Supp. Info. 2), implemented support for Cyanogate assemblies in the online DNA “Design and Build” portal of the Edinburgh Genome Foundry (dab.genomefoundry.org) (Supp. Info. 9), and submitted all vectors as a toolkit for order from Addgene (www.addgene.org).

Standardisation will help to accelerate the development of reliable synthetic biology tools for biotechnological applications and promote sharing and evaluation of genetic parts in different species and under different culturing conditions (40). Going forward, it will be important to test the performance of different parts with different components (e.g. gene expression cassettes) and in different assembly combinations. Several groups using plant MoClo assembly have reported differences in cassette expression and functionality depending on position and orientations (e.g. 92), which highlights a key synthetic biology crux - the performance of a system is not simply the sum of its components (93, 94).

The increasing availability of genome-scale metabolic models for different cyanobacterial species and their utilisation for guiding engineering strategies for producing heterologous high-value biochemicals has helped to re-invigorate interest in the industrial potential of cyanobacteria (95-98). Future efforts should focus on combining genome-scale metabolic models with synthetic biology approaches, which may help to overcome the production yield limitations observed for cyanobacterial cell factories (8), and will accelerate the development of more complex and precise gene control circuit systems that can better integrate with host metabolism and generate more robust strains (99-101). The future development of truly ‘programmable’ photosynthetic cells could provide significant advancements in addressing fundamental biological questions and tackling global challenges, including health and food security (102-104).

## SUPPLEMENTARY DATA

**Supporting Information 1. Table of all parts from CyanoGate kit generated in this work**. Domestication refers to the removal of *BsaI* and/or *BsiI* sites (modifications are indicated in sequence maps provided in **Supp. Info. 2**).

**Supporting Information 2. Sequence maps (.gb files) of the components of the CyanoGate kit**.

**Supporting Information 3. Protocols for MoClo assembly in level -1 through to level T**. Protocols for assembly in level 0, level M and level T acceptor vectors (restriction enzyme *Bpi*I required, left). Protocols for assembly in level -1, level 1 and level P backbone vectors (restriction enzyme *Bsa*I required, right). Adapted from “A quick guide to Type IIS cloning” (Patron Lab; http://patronlab.org/). For troubleshooting Type IIS mediated assembly we recommend http://synbio.tsl.ac.uk/docs.

**Supporting Information 4. Detailed assembly strategies using the CyanoGate kit**.

**Supporting Information 5. Integrative engineering strategies using the CyanoGate kit**. (A) Marked mutants are generated using a level T marked knock out vector carrying DNA sequences flanking the target locus of the chromosome (~1 kb), an antibiotic resistance cassette (Ab*^R^*) and a sucrose selection cassette (*sac*B) that produces the toxic compound levansucrase in the presence of sucrose (20). Several rounds of segregation are required to identify a marked mutant. (B) Marked mutants then can be unmarked with a level T unmarked knock out vector and selection on sucrose-containing agar plates. (C) Unmarked knock in mutants can also be generated from marked mutants using a level T unmarked knock in vector carrying a gene expression cassette (UP FLANK LINKER and DOWN FLANK LINKER are shown in pink and light green, respectively). (D) Alternatively marked knock in mutants can be engineered in a single step using a level T marked knock vector (Ab^R^ UP LINKER and DOWN LINKER are shown in blue and orange, respectively). See **Fig. 2** for abbreviations.

**Supporting Information 6. Comparison of growth for *Synechocystis*, PCC 7942 and UTEX 2973 under different culturing conditions**. Values are the means ± SE from at least five biological replicates from two independent experiments.

**Supporting Information 7. Growth and expression levels of heterologous and synthetic promoters in *Synechocystis* and UTEX 2973**. (A) *Synechocystis* and UTEX 2973 was cultured for 72 hr at 30°C with continuous light (100 μmol photons m^−2^ s^−1^) and 40°C with 300 μmol photons m^−2^ s^−1^, respectively (see **Fig 6**). Expression levels of eYFP are shown at three time points. Values are the means ± SE from at least four biological replicates.

**Supporting Information 8. List of level T vectors used in this study**.

**Supporting Information 9. Protocol and online interface for building CyanoGate vector assemblies**. A CyanoGate online vector assembly tool called Design and Build (DAB) from the Edinburgh Genome Foundry.

## ACKNOWLEDGEMENTS

We thank Conrad Mullineaux (Queen Mary University), Julie Zedler and Poul Erik Jensen (University of Copenhagen) for providing components for conjugation, and Eva Steel (University of Edinburgh) for assistance with assembling the CyanoGate kit.

## References

1. Castenholz, R.W. (2001) Phylum BX. Cyanobacteria. Oxygenic photosynthetic bacteria. In: Garrity, G., Boone, D.R., Castenholz, R.W. (eds) Bergey’s manual of systematic bacteriology. Springer, New York, pp 473–599.

2. Flombaum, P., Gallegos, J.L., Gordillo R.A., Rincón, J., Zabala, L.L., Jiao, N., Karl, D.M., Li, W.K.W., Lomas, M.W., Veneziano, D., Vera, C.S., Vrugt, J.A., Martiny, A.C. (2013) Present and future global distributions of the marine Cyanobacteria Prochlorococcus and Synechococcus. Proc. Natl. Acad. Sci. U.S.A., 110, 9824–9829.

3. Keeling, P.J. (2003) Diversity and evolutionary history of plastids and their hosts. Am. J. Bot., 10, 1481–1493.

4. Rae, B.D., Long, B., Förster, B., Nguyen, N.D., Velanis, C.N., Atkinson, N., Hee, W., Mukherjee, B., Price, G.D., McCormick, A.J. (2017) Progress and challenges of engineering a biophysical carbon dioxide concentrating mechanism into higher plants. J. Ex. Bot., 68, 3717–3737.

5. Tan, X., Yao, L., Gao, Q., Wang, W., Qi, F., Lu, X. (2011) Photosynthesis driven conversion of carbon dioxide to fatty alcohols and hydrocarbons in cyanobacteria. Metab. Eng., 13, 169–176.

6. Ducat, D.C., Way, J.C., Silver, P.A. (2010) Engineering cyanobacteria to generate high-value products. Trends Biotechnol., 29, 95–103.

7. Ramey, C. J., Barón-Sola, Á., Aucoin, H.R., Boyle, N.R. (2015) Genome Engineering in Cyanobacteria: Where We Are and Where We Need To Go. ACS Synth. Biol., 4, 1186–1196.

8. Nielsen, A.Z., Mellor, S.B., Vavitsas, K., Wlodarczyk, A.J., Gnanasekaran, T., de Jesus, M.P.R.H., King, B.C. Bakowski, K., Jensen, P.E. (2016) Extending the biosynthetic repertoires of cyanobacteria and chloroplasts. Plant J., 87, 87–102.

9. Wlodarczyk, A., Gnanasekaran, T., Nielsen, A.Z., Zulu, N.N., Mellor, S.B., Luckner, M., Thøfner, J.F.B, Olsen, C.E., Mottawie, M.S. Burow, M. Pribil, M. Feussner, I., Møller, B.L. Jensen, P.E. (2016) Metabolic engineering of light-driven cytochrome P450 dependent pathways into *Synechocystis* sp. PCC 6803. Metab. Eng., 33, 1–11.

10. Pye, C.R., Bertin, M.J., Lokey, R.S., Gerwick, W.H., Linington, R.G. (2017) Retrospective analysis of natural products provides insights for future discovery trends. Proc. Natl. Acad. Sci. U.S.A., 114, 5601–5606.

11. Stensjö, K., Vavitsas, K., Tyystjärvi, T. (2017) Harnessing transcription for bioproduction in cyanobacteria. Physiol. Plant, 162, 148–155.

12. McCormick, A.J., Bombelli, P., Bradley, R.W., Thorne, R., Wenzele, T., Howe, C.J. (2015) Biophotovoltaics: oxygenic photosynthetic organisms in the world of bioelectrochemical systems. Energy Environ. Sci., 8, 1092–1109.

13. Saar, K.L., Bombelli, P., Lea-Smith, D.J., Call, T., Aro, E-M., Müller, T., Howe, C.J., Knowles, T.P.J (2018) Enhancing power density of biophotovoltaics by decoupling storage and power delivery. Nat. Energy, 3, 75–81.

14. Vioque, A. (2007) Transformation of Cyanobacteria. In: León, R., Galván, A., Fernández, E. (eds) Transgenic Microalgae as Green Cell Factories. Advances in Experimental Medicine and Biology, vol 616. Springer, New York, pp 12–22.

15. Stucken, K., Ilhan, J., Roettger, M., Dagan, T. Martin, W.F. (2012) Transformation and conjugal transfer of foreign genes into the filamentous multicellular cyanobacteria (subsection V) *Fischerella* and *Chlorogloeopsis*. Curr. Microbiol., 65, 552–560.

16. Heidorn, T., Camsund, D., Huang, H., Lindberg, P., Oliveira, P., Stensjö, K., Lindblad, P. (2011) Synthetic biology in cyanobacteria engineering and analyzing novel functions. Methods Enzymol., 497, 539–579.

17. Mermet-Bouvier, P., Cassier-Chauvat, C., Marraccini, P., Chauvat, F. (1993) Transfer and replication of RSF1010-derived plasmids in several cyanobacteria of the general *Synechocystis* and *Synechococcus*. Curr. Microbiol. 27, 323–327.

18. Huang, H., Camsund, D., Lindblad, P., Heidorn, T. (2010) Design and characterization of molecular tools for a Synthetic Biology approach towards developing cyanobacterial biotechnology. Nucleic Acids Res., 38, 2577–2593.

19. Taton, A.U., Unglaub, F. Wright, N.E., Zeng, W.Y., Paz-Yepes, J., Brahamsha, B., Palenik, B. Peterson, T.C., Haerizadeh, F., Golden, S.S., Golden, J.W. (2014) Broad-host-range vector system for synthetic biology and biotechnology in cyanobacteria. Nucleic Acids Res., 42, e136.

20. Lea-Smith, D.J., Vasudevan, R., Howe, C.J. (2016) Generation of marked and markerless mutants in model cyanobacterial species. J. Vis. Exp., 111, e54001.

21. Ungerer, J., Pakrasi, H.B. (2016) Cpf1 Is a versatile tool for CRISPR genome editing across diverse species of cyanobacteria. Sci. Rep., 6, e39681.

22. Wendt, K.E., Ungerer, J., Cobb, R.E., Zhao, H., Pakrasi, H.B. (2016) CRISPR/Cas9 mediated targeted mutagenesis of the fast growing cyanobacterium *Synechococcus elongatus* UTEX 2973. Microb. Cell. Fact., 15, e115

23. Huang, H., Lindblad, P. (2013) Wide-dynamic-range promoters engineered for cyanobacteria. J. Biol. Eng., 7, 10.

24. Camsund, D., Heidorn, T., and Lindblad, P. (2014) Design and analysis of LacI-repressed promoters and DNA-looping in a cyanobacterium. J. Biol. Eng., 8, e4.

25. Albers, S.C., Gallegos, V.A., Peebles, C.A.M. (2015) Engineering of genetic control tools in *Synechocystis* sp. PCC 6803 using rational design techniques. J. Biotechnol., 216, 36–46.

26. Markley, A.L., Begemann, M.B., Clarke, R.E., Gordon, G.C., Pflege, B.F. (2015) Synthetic biology toolbox for controlling gene expression in the cyanobacterium *Synechococcus* sp. strain PCC 7002. ACS Synth Biol., 4, 595–603.

27. Englund, E., Liang, F., Lindberg, P. (2016) Evaluation of promoters and ribosome binding sites for biotechnological applications in the unicellular cyanobacterium *Synechocystis* sp. PCC 6803. Sci. Rep., 6, A36640.

28. Kim, W.J., Lee, S.M., Um, Y., Sim, S.J., Woo, H.M. (2017) Development of SyneBrick vectors as a synthetic biology platform for gene expression in *Synechococcus elongatus* PCC 7942. Front. Plant Sci., 8, A293.

29. Immethun, C.M., DeLorenzo, D.M., Focht, C.M., Gupta, D., Johnson, C.B., Moon, T.S. (2017) Physical, chemical, and metabolic state sensors expand the synthetic biology toolbox for *Synechocystis* sp. PCC 6803. Biotechnol Bioeng., 114, 1561–1569.

30. Taton, A. U., Ma, A.T., Ota, M., Golden, S.S., Golden, J.W. (2017) NOT gate genetic circuits to control gene expression in cyanobacteria. ACS Synth. Biol., 6, 2175–2182.

31. Ferreira, E.A., Pacheco, C.C., Pinot, F., Pereira, J., Lamosa, P., Oliveira, P., Kirov, B., Jaramillo, A., Tamagnini, P. (2018) Expanding the toolbox for Synechocystis sp. PCC 6803: validation of replicative vectors and characterization of a novel set of promoters. Synth. Biol., 3, ysy014.

32. Li, S., Sun, T., Xu, C., Chen, L., Zhang, W. (2018) Development and optimization of genetic toolboxes for a fast-growing cyanobacterium *Synechococcus elongatus* UTEX 2973. Metab. Eng., 48, 163–174.

33. Lui, D., Pakrasi, B.H. (2018) Exploring native genetic elements as plug-in tools for synthetic biology in the cyanobacterium *Synechocystis* sp. PCC 6803. Microb. Cell Fact., 17, 48.

34. Wang, B., Eckert, C., Maness, P-C., Yu, J. (2018) A genetic toolbox for modulating the expression of heterologous genes in the cyanobacterium *Synechocystis* sp. PCC 6803. ACS Synth. Biol., 19, 276–286.

35. Nielsen, J., Keasling, J.D. (2016) Engineering cellular metabolism. Cell, 164, 1185–1197.

36. Moore, S.J., Lai, H-E., Kelwick, R.J.R., Chee, S.M., Bell, D.J., Polizzi, K.M., Freemont, P.S. (2016) EcoFlex: a multifunctional MoClo kit for *E. coli* synthetic biology. ACS Synth. Biol., 5, 1059–1069.

37. Sarrion-Perdigones, A., Vazquez-Vilar, M., Palací, J., Castelijns, B., Forment, J., Ziarsolo, P., Blanca, J., Granell, A., Orzaez, D. (2013) GoldenBraid 2.0: a comprehensive DNA assembly framework for plant synthetic biology. Plant Physiol., 162, 1618–1631.

38. Engler, C., Youles, M., Gruetzner, R., Ehnert, T-M., Werner, S., Jones, J.D.G., Patron, N.J., Marillonnet, S. (2014) A Golden Gate modular cloning toolbox for plants. ACS Synth. Biol., 3, 839–843.

39. Andreou, A.I., Nakayama, N. (2018) Mobius Assembly: A versatile Golden-Gate framework towards universal DNA assembly. PLoS ONE, 13, e0189892.

40. Patron, N.J., Orzaez, D., Marillonnet, S., Warzecha, H., Matthewman, C., Youles, M., Raitskin, O., Leveau, A., Farré, G., Rogers, C., Smith, A., Hibberd, J., Webb, A.A. et al. (2015) Standards for plant synthetic biology: a common syntax for exchange of DNA parts. New Phytol., 208, 13–19.

41. Chambers, S., Kitney, R., Freemont, P. (2016) The Foundry: the DNA synthesis and construction Foundry at Imperial College. Biochem. Soc. Trans., 44, 687–688.

42. Crozet, P., Navarro, F.J., Willmund, F., Mehrshahi, P., Bakowski, K., Lauersen, K.J., Pérez-Pérez, M.E., Auroy, P., Gorchs Rovira, A., Sauret-Gueto, S., Niemeyer, J., Spaniol, B., Theis, J., Trösch, R., Westrich, L.D., Vavitsas, K., Baier, T., Hübner, W., de Carpentier, F., Cassarini, M., Danon, A., Henri, J., Marchand, C.H., de Mia, M., Sarkissian, K., Baulcombe, D.C., Peltier, G., Crespo, J.L., Kruse, O., Jensen, P.E., Schroda, M., Smith, A.G., Lemaire, S.D. (2018) Birth of a photosynthetic chassis: A MoClo toolkit enabling synthetic biology in the microalga *Chlamydomonas reinhardtii*. ACS Synth. Biol., 7, 2074–2086.

43. Na, D., Yoo, S.M., Chung, H., Park, H., Park, J.H., Lee, S.Y. (2013) Metabolic engineering of Escherichia coli using synthetic small regulatory RNAs. Nat. Biotechnol., 31, 170–174.

44. Yao, L., Cengic, I., Anfelt, J., Hudson, E.P. (2015) Multiple gene repression in cyanobacteria using CRISPRi. ACS Synth. Biol., 5, 207–212.

45. Sun, T., Li, S., Song, X., Diao, J., Chen, L., Zhang, W. (2018) Toolboxes for cyanobacteria: Recent advances and future direction. Biotechnol Adv., 36, 1293–1307.

46. Yu, J., Liberton, M., Cliften, P.F., Head, R.D., Jacobs, J.M., Smith, R.D, Koppenaal, D.W., Brand, J.J., Pakrasi, H.B. (2015) *Synechococcus elongatus* UTEX 2973, a fast growing cyanobacterial chassis for biosynthesis using light and CO_2_. Sci. Rep., 5, 8132.

47. Ungerer, J., Lin, P-C., Chen, H-Y., Pakrasi, H.B. (2018) Adjustments to photosystem stoichiometry and electron transfer proteins are key to the remarkably fast growth of the cyanobacterium *Synechococcus elongatus* UTEX 2973. mBio, 9, e02327–17.

48. Rippka, R., Deruelles, J., Waterbury, J.B., Herdman, M., Stanier, R.Y. (1979) Generic assignments, strain histories and properties of pure cultures of cyanobacteria. J. Gen. Microbiol., 111, 1–61.

49. Kelly, C.L., Hitchcock, A., Torres-Méndez, A., Heap, J.T. (2018) A rhamnose-inducible system for precise and temporal control of gene expression in cyanobacteria. ACS Synth. Biol., 7, 1056–1066.

50. Silva-Rocha, R., Martínez-García, E., Calles, B., Chavarría, M., Arce-Rodríguez, A., de Las Heras, A., Páez-Espino, A.D., Durante-Rodríguez, G., Kim, J., Nikel, P.I., Platero, R., de Lorenzo, V (2013) The Standard European Vector Architecture (SEVA): a coherent platform for the analysis and deployment of complex prokaryotic phenotypes. Nucleic Acids Res. 41, D666–675.

51. Liu, Q., Schumacher, J., Wan, X., Lou, C., Wang, B. (2018) Orthogonality and burdens of heterologous AND Gate gene circuits in E. coli. ACS Synth. Biol., 7, 553–564.

52. Qi, Q., Hao, M., Ng, W., Slater, S.C., Baszis, S.R., Weiss, J.D., Valentin, H.E. (2005) Application of the *Synechococcus nirA* promoter to establish an inducible expression system for engineering the *Synechocystis* tocopherol pathway. Appl. Environ. Microbiol., 71, 5678–5684.

53. Tsinoremas, N.F., Kutach, A.K., Strayer, C.A., Golden, S.S. (1994) Efficient gene transfer in *Synechococcus* sp. strains PCC 7942 and PCC 6301 by interspecies conjugation and chromosomal recombination. J. Bacteriol., 176, 6764–6768.

54. Werner, S., Engler, C., Weber, E., Gruetzner, R., Marillonnet, S. (2012) Fast track assembly of multigene constructs using Golden Gate cloning and the MoClo system. Bioeng. Bugs. 3, 38–43.

55. Kirst, H., Formighieri, C., Melis, A. (2014) Maximizing photosynthetic efficiency and culture productivity in cyanobacteria upon minimizing the phycobilisome light-harvesting antenna size. Biochim. Biophys. Acta, 1837, 1653–1664.

56. Lea-Smith, D.J., Bombelli, P., Dennis, J.S., Scott, S.A., Smith, A.G. Howe, C.J. (2014) Phycobilisome-deficient strains of *Synechocystis* sp. PCC 6803 have reduced size and require carbon-limiting conditions to exhibit enhanced productivity. Plant Physiol., 165, 705–714.

57. Liberton, M., Chrisler, W.B., Nicora, C.D., Moore, R.J., Smith, R. D., Koppenaal, D.W., Pakrasi, H.B., Jacobs, J.M. (2017) Phycobilisome truncation causes widespread proteome changes in *Synechocystis* sp. PCC 6803. PLoS ONE, 12, e0173251

58. Pinto, F., Pacheco, C. C., Oliveira, P., Montagud, A., Landels, A., Couto, N., Wright, P. C., Urchueguía, J. F., Tamagnini, P. (2015) Improving a Synechocystis-based photoautotrophic chassis through systematic genome mapping and validation of neutral sites. DNA Res., 22, 425–437.

59. Vogel, A.I.M, Lale, R. Hohmann-Marriott, M.F. (2017) Streamlining recombination-mediated genetic engineering by validating three neutral integration sites in *Synechococcus* sp. PCC 7002. J. Biol. Eng., 11, 19.

60. Kulkarni, D.R., Golden S.S. (1997). mRNA stability is regulated by a coding-region element and the unique 5’ untranslated leader sequences of the three Synechococcus psbA transcripts. Mol. Microbiol., 24, 1131–1142.

61. Andersson, C.R., Tsinoremas, N.F., Shelton, J., Lebedeva, N.V., Yarrow, J., Min, H., Golden, S.S. (2000) Application of bioluminescence to the study of circadian rhythms in cyanobacteria. Methods Enzymol., 305, 527–542.

62. Chang, Y.G., Cohen, S.E., Phong, C., Myers, W.K., Kim, Y.I., Tseng, R., Lin, J., Zhang, L., Boyd, J.S., Lee, Y., Kang, S., Lee, D., Li, S., Britt, R.D., Rust, M.J., Golden, S.S., LiWang, A. (2015) Circadian rhythms. A protein fold switch joins the circadian oscillator to clock output in cyanobacteria. Science, 349, 324–328.

63. Peca, L., Kós, P.B., Máté, Z., Farsang, A., Vass, I. (2008) Construction of bioluminescent cyanobacterial reporter strains for detection of nickel, cobalt and zinc. FEMS Microbiol. Lett., 289, 264.

64. Blasi, B., Peca, L., Vass, I., Kós, P.B. (2012) Characterization of stress responses of heavy metal and metalloid inducible promoters in *Synechocystis* PCC6803. J. Microbiol. Biotechnol., 22, 166–169.

65. Guerrero, F., Carbonell, V., Cossu, M., Correddu, D., Jones, P.R. (2012) Ethylene synthesis and regulated expression of recombinant protein in *Synechocystis* sp. PCC 6803. PLoS ONE, 7, e50470.

66. Kikuchi, H., Aichi, M., Suzuki, I., Omato, T. (1996) Positive regulation by nitrite of the nitrate assimilation operon in the cyanobacteria *Synechococcus* sp. strain PCC 7942 and *Plectonema boryanum*. J. Bacteriol., 178, 5822–5825.

67. Kunert, A., Vinnemeier, J., Erdmann, N., Hagemann, M. (2003) Repression by Fur is not the main mechanism controlling the iron-inducible isiAB operon in the cyanobacterium *Synechocystis* sp. PCC 6803. FEMS Microbiol. Lett., 227, 255–262.

68. Lindberg, P., Park, S., Melis, A. (2010) Engineering a platform for photosynthetic isoprene production in cyanobacteria, using *Synechocystis* as the model organism. Metab. Eng., 12, 70–79.

69. Zhou, J., Zhang, H., Meng, H., Zhu, H., Bao, G., Zhang, Y., Li, Y., Ma, Y. (2014) Discovery of a super-strong promoter enables efficient production of heterologous proteins in cyanobacteria. Sci. Rep., 4, 4500.

70. Munch, R., Hiller, K., Grote, A., Scheer, M., Klein, J., Schobert, M., Jahn, D. (2005) Genome analysis Virtual Footprint and PRODORIC: an integrative framework for regulon prediction in prokaryotes. Bioinforma. Appl. Note 21, 4187–4189.

71. Bhal, C.P., Wu, R., Stawinsky, J. Narang, S.A. (1977) Minimal length of the lactose operator sequence for the specific recognition by the lactose repressor. Proc. Natl. Acad. Sci. U.S.A., 74, 966–970.

72. Abe, K., Sakai, Y., Nakashima, S., Araki, M., Yoshida, W., Sode, K., Ikebukuro, K. (2014) Design of riboregulators for control of cyanobacterial (*Synechocystis)* protein expression. Biotechnol. Lett., 36, 287–294.

73. Ng, A.H., Berla, B.M., Pakrasi, H.B. Fine-tuning of photoautotrophic protein production by combining promoters and neutral sites in the cyanobacterium *Synechocystis* sp. strain PCC 6803. Appl. Environ. Microbiol. 81, 6857–6863.

74. van de Meene, A.M., Hohmann-Marriott, M.F., Vermaas, W.F., Roberson, R.W. (2006) The three-dimensional structure of the cyanobacterium *Synechocystis* sp. PCC 6803. Arch Microbiol., 184, 270.

75. Mueller, T.J., Ungerer, J.L., Pakrasi, H.B., Maranas, C.D. (2017) Identifying the metabolic differences of a fast-growth phenotype in Synechococcus UTEX 2973. Sci. Rep., 7, 41569.

76. Bradley, R.W., Buck, M., Wang, B. (2016) Tools and principles for microbial gene circuit engineering. J. Mol. Biol., 428, 862–882.

77. Armshaw, P., Carey, D., Sheahan, C., Pembroke, J.T. (2015) Utilising the native plasmid, pCA2.4, from the cyanobacterium *Synechocystis* sp. strain PCC6803 as a cloning site for enhanced product production. Biotechnol. Biofuels 8, 201.

78. Chen, Y., Taton, A., Go, M., London, R.E., Pieper, L.M., Golden, S.S., Golden, J.W. (2016). Self-replicating shuttle vectors based on pANS, a small endogenous plasmid of the unicellular cyanobacterium *Synechococcus elongatus* PCC 7942. Microbiology, 162, 2029–2041.

79. Frey, J., Bagdasarian, M.M., Bagdasarian, M. (1992) Replication and copy number control of the broad-host-range plasmid RSF1010. Gene, 113, 101–106.

80. Blasina, A., Kittell, B.L., Toukdarian, A.E., Helinski, D.R. (1996) Copy-up mutants of the plasmid RK2 replication initiation protein are defective in coupling RK2 replication origins. Proc. Natl. Acad. Sci. U.S.A., 93, 3559–3564.

81. Qi, L.S., Larson, M.H., Gilbert, L.A., Doudna, J.A., Weissman, J.S. Arkin, A.P., Lim, W.A. (2013) Repurposing CRISPR as an RNA-guided platform for sequence-specific control of gene expression. Cell, 152, 1173–1183.

82. Behler, J., Vijay, D., Hess, W.R., Akhtar, M.K. (2018) CRISPR-based technologies for metabolic engineering in cyanobacteria. Trends Biotechnol., pii: S0167-7799(18)30146-X.

83. Huang, C.H., Shen, C.R., Li, H., Sung, L.Y., Wu, M.Y., Hu, Y.C. (2016) CRISPR interference (CRISPRi) for gene regulation and succinate production in cyanobacterium S. elongatus PCC 7942. Microb. Cell Fact., 15, 196.

84. Higo, A., Isu, A., Fukaya, Y., Ehira, S., Hisabori, T. (2018) Application of CRISPR interference for metabolic engineering of the heterocyst-forming multicellular cyanobacterium Anabaena sp. PCC 7120. Plant Cell Physiol., 59, 119–127.

85. Zess, E.K., Begemann, M.B., Pfleger, B.F. (2016) Construction of new synthetic biology tools for the control of gene expression in the cyanobacterium Synechococcus sp. strain PCC 7002. Biotechnol. Bioeng., 113, 424–432.

86. Sakai, Y., Abe, K., Nakashima, S., Ellinger, J.J., Ferri, S., Sode, K., Ikebukuro, K. (2015) Scaffold-fused riboregulators for enhanced gene activation in *Synechocystis* sp. PCC 6803. MicrobiologyOpen, 4, 533–540.

87. Xiao, A., Cheng, Z., Kong, L., Zhu, Z., Lin, S., Gao, G., Zhang, B. (2014) CasOT: a genome-wide Cas9/gRNA off-target searching tool. Bioinformatics, 30, 1180–1182.

88. Gordon, G.C., Korosh, T.C., Cameron, J.C., Markley, A.L., Begemann, M.B., Pfleger, B.F. (2016) CRISPR interference as a titratable, trans-acting regulatory tool for metabolic engineering in the cyanobacterium *Synechococcus* sp. strain PCC 7002. Metab. Eng., 38, 170–179.

89. Economou, C., Wannathong, T., Szaub, J., Purton, S. (2014) A simple, low-cost method for chloroplast transformation of the green alga *Chlamydomonas reinhardtii*. Methods Mol. Biol., 1132, 401–411.

90. Doud, D.F.R., Holmes, E.C., Richter, H., Molitor, B., Jander, G., Angenent, L.T. (2017) Metabolic engineering of Rhodopseudomonas palustris for the obligate reduction of n-butyrate to n-butanol. Biotechnol. Biofuels., 10, 178.

91. Leonard, S.P., Perutka, J., Powell, J.E., Geng, P., Richhart, D.D., Byrom, M., Kar, S., Davies, B.W., Ellington, A.D., Moran, N.A., Barrick, J.E. (2018) Genetic engineering of bee gut microbiome bacteria with a toolkit for modular assembly of broad-host-range plasmids. ACS Synth. Biol., 7, 1279–1290.

92. Ordon, J., Gantner, J., Kemna, J., Schwalgun, L., Reschke, M., Streubel, J., Boch, J., Stuttmann, J. (2017) Generation of chromosomal deletions in dicotyledonous plants employing a user-friendly genome editing toolkit. Plant J., 89:155–168.

93. Mutalik, V.K., Guimaraes, J.C., Cambray, G., Mai, Q.A., Christoffersen, M.J., Martin, L., Yu, A., Lam, C., Rodriguez, C., Bennett, G., Keasling, J.D., Endy, D., Arkin, A.P. (2013) Quantitative estimation of activity and quality for collections of functional genetic elements. Nat. Methods 10, 347–353.

94. Heyduk, E., Heyduk, T. (2018) DNA template sequence control of bacterial RNA polymerase escape from the promoter. Nucleic Acids Res., 46, 4469–4486.

95. Knoop, H., Gründel, M., Zilliges, Y., Lehmann, R., Hoffmann, S., Lockau, W., Steuer, R. (2013) Flux Balance Analysis of Cyanobacterial Metabolism: The Metabolic Network of *Synechocystis* sp. PCC 6803. PLoS Comput. Biol., 9, e1003081.

96. Hendry, J.I., Prasannan, C.B., Joshi, A., Dasgupta, S., Wangikar, P.P. (2016) Metabolic model of *Synechococcus* sp. PCC 7002: Prediction of flux distribution and network modification for enhanced biofuel production. Bioresour Technol., 213, 190–197.

97. Mohammadi, R., Fallah-Mehrabadi, J., Bidkhori, G., Zahiri, J., Javad Niroomand, M., Masoudi-Nejad, A. (2016) A systems biology approach to reconcile metabolic network models with application to Synechocystis sp. PCC 6803 for biofuel production. Mol. Biosyst., 12, 2552–2561.

98. Shirai, T., Osanai, T., Kondo, A. (2016) Designing intracellular metabolism for production of target compounds by introducing a heterologous metabolic reaction based on a Synechosystis sp. 6803 genome-scale model. Microb. Cell Fact., 15, 13.

99. Bradley, R.W., Wang, B. (2015) Designer cell signal processing circuits for biotechnology. N. Biotechnol., 32, 635–643.

100. Jusiak, B., Cleto, S., Perez-Piñera, P., Lu, T.K. (2016) Engineering synthetic gene circuits in living cells with CRISPR technology. Trends Biotechnol., 34, 535–547.

101. Luan, G., Lu, X. (2018) Tailoring cyanobacterial cell factory for improved industrial properties. Biotechnol. Adv., 36, 430–442.

102. Dobrin, A., Saxena, P., Fussenegger, M. (2016) Synthetic biology: applying biological circuits beyond novel therapies. Integr. Biol., 8, 409–430.

103. Medford. J.I., Prasad. A. (2016) Towards programmable plant genetic circuits. Plant J., 87, 139–148.

104. Smanski, M.J., Zhou, H., Claesen, J., Shen, B., Fischbach, M.A., Voigt, C.A. (2016) Synthetic biology to access and expand nature’s chemical diversity. Nat. Rev. Microbiol., 14, 135–149.

